# Influenza A Virus Infection Induces Immune Dysregulation in the Placenta and Fetus Without Vertical Transmission in Nonhuman Primates

**DOI:** 10.1101/2025.10.06.675688

**Authors:** Orlando Cervantes, Sidney Sun, Hazel Huang, Briana Del Rosario, Hong Zhao, Amanda Li, Andrew Vo, Gygeria Manuel, Roslyn Van Abel, Jeff Munson, John Cornelius, Raj P. Kapur, Miranda Li, Edmunda Li, Hanning Li, Alexandra Christodoulou, Melissa Berg, Britni Curtis, Elizabeth Miller, Shannon Ruff, Brenna Menz, Solomon Wangari, Audrey Germond, Audrey Baldessari, Chris English, W. McIntyre Durning, Thomas B. Lewis, Megan N. Fredericks, Mara Maughan, Austyn Orvis, Michelle Coleman, Lakshmi Rajagopal, Deborah H. Fuller, Kristina M. Adams Waldorf

## Abstract

Influenza A virus (IAV) infection during pregnancy is associated with stillbirth and preterm birth, but the degree to which IAV alters placental and fetal immunity is poorly understood. The objective of our study was to determine the immunologic impact of maternal IAV infection on the placenta and fetus in a pigtail macaque (*Macaca nemestrina*) model. Pregnant pigtail macaques were inoculated with 10^7^ plaque forming units (PFU) of IAV [A/California/07/2009 (H1N1)] and underwent necropsy 5 days post-infection (N=11). Results were compared to uninfected historical controls (N=16). IAV inoculation induced maternal pneumonia in all cases. Stillbirth occurred in 18% (2/11) of IAV-infected pregnancies, but not in controls. While vertical transmission was not observed, low-level IAV viral RNA was detected in two placentas. In the placenta, maternal IAV infection was associated with increased IL-1β, IL-18, and IFN-β levels, and an upregulated type I interferon (IFN) transcriptional response. IAV infection was also associated with significantly higher frequencies of intermediate and non-classical monocytes, plasmacytoid dendritic cells, CD4⁺ T cells, and NKT cells in the fetus (lung, lymph node, blood). Although placental immune and transcriptional perturbations were rarely correlated with maternal IAV disease indicators (e.g., maternal lung viral load/IFN-α/IFN-β/IL-6), there were consistent and significant correlations between these metrics and perturbed immune cell populations in the fetus (CD4+ and CD8+ T cells, plasmacytoid dendritic cells, monocyte sub-populations). Maternal IAV infection disrupted both placental and fetal immune environments, but only fetal immune alterations correlated with maternal lung disease severity.

**One Sentence Summary:** Maternal influenza A virus infection in pregnant pigtail macaques dysregulates placental and fetal immunity, with disease severity correlating strongly with fetal, but not placental, immune perturbations.

## INTRODUCTION

Influenza A virus (IAV) infections remain a global public health threat, which is especially pronounced in immunologically vulnerable populations like pregnant women.^1,2^ IAV is a negative-sense, single-stranded RNA virus composed of eight segments. Pandemics occur when reassortment of viral segments produces new combinations of its surface glycoproteins, hemagglutinin (HA) and neuraminidase (NA). Epidemiologic evidence from several IAV pandemics (e.g., 1918, 1957, 1968, 2009) indicates that IAV infections in pregnancy impart a greater risk of maternal mortality and adverse pregnancy outcomes, such as stillbirth.(*1-12*) There may also be a long-term risk of neuropsychiatric disease in the children born to pregnant women infected with IAV or other infectious diseases, which has been reported in multiple populations.(*13-20*)

The effects of placental and fetal exposure to maternal influenza A virus (IAV) have been studied primarily in murine models. Experimental IAV infection in pregnant mice and rats (e.g., H1N1, H3N2) has been associated with a range of adverse outcomes, including preterm birth, low birth weight, and stillbirth (resorptions).(*21-25*) These models demonstrate robust activation of innate and adaptive immunity in the maternal compartment, with increased monocytes, neutrophils, and T cells detected in the lung, placenta, and gut-associated lymphoid tissue. In mice, IAV can disrupt progesterone and prostaglandin signaling in the placenta, which was associated with adverse perinatal outcomes and placental remodeling.(*24, 26*) Severe maternal IAV-associated vascular inflammation has been associated with placental growth restriction and fetal brain hypoxia.(*21*) In contrast, relatively few studies have examined fetal immune perturbations following in utero exposure to maternal IAV, with reports describing only altered fetal thymic transcription and reduced neutralizing antibody responses to IAV.(*22, 23*) Thus, the extent to which systemic maternal disease coordinates immune changes across the placenta and fetus, whether these responses scale with disease severity, and how they ultimately influence fetal development remain unclear.

The objective of our study was to determine the immunologic impact of a maternal infection with an IAV H1N1 2009 pandemic strain on the placenta and fetus in a pigtail macaque (*Macaca nemestrina*) model, with a focus on whether severity of maternal lung disease influenced these changes. While some of these molecular mechanisms have been identified in mice and rats, these animal models differ from human pregnancy in many respects, including a short gestation (3-5 weeks). Humans have a gestational period that is 8–10 times longer than these species, allowing for extended fetal brain development *in utero.* However, this long gestational period also increases placental and fetal exposure to seasonal waves of IAV infection. Nonhuman primates (NHPs) share many similarities with human pregnancy, including longer gestation, placental structure, and labor physiology.(*27, 28*) NHPs are also frequently used in IAV disease modeling due to their similarities in human clinical and pathological features.(*29, 30*) A few studies using pregnant NHPs have modeled IAV infection but were performed to specifically understand complement activation and infant brain development in African green monkeys and rhesus macaques, respectively.(*31, 32*) Thus, our central goal was to broadly characterize how IAV infection during pregnancy affects placental and fetal immunity in a pigtail macaque model, a highly translational model to human pregnancy.

## RESULTS

### Impact of Maternal IAV Disease on Clinical Outcomes and Histopathology

Eleven pregnant pigtail macaques were inoculated with an IAV H1N1 2009 pandemic strain (A/CA/07/2009) in the third trimester (FLU cohort) using a combination of intranasal, intratracheal, intraocular, and intraoral routes with a total of 10^7.4^ plaque-forming units (PFU). Results were compared to 16 historical control uninfected pregnant pigtail macaques (CTRL cohort, Table S1), which received either subcutaneous media inoculations (N=9)(*33*) or saline inoculation into the choriodecidual space via catheters (N=7)(*34*) for comparison against Zika virus or Group B *Streptococcus* cohorts, respectively. Five days after inoculation, a Cesarean section and necropsies of the dam and fetus were performed. The mean gestational age on the day of Cesarean section was 133.4 days in the FLU cohort, 136.7 days in the CTRL catheterized group, and 153.1 days in the CTRL media control cohort

Clinical signs of maternal illness were documented using daily clinical scores and vital signs taken at the time of sedations: before inoculation, 1- and 3 days post-inoculation (dpi), and on 5 dpi, the experimental endpoint (Table 1). Coughing, wheezing, increased respiratory effort, nasal discharge, and decreased appetite were common symptoms in 7 of 11 FLU animals (64%), while the remaining four did not show clinical symptoms. Animals that exhibited the most severe disease based on clinical scores were FLU8 and FLU9; of these animals, FLU9 experienced a stillbirth. Clinical scores were highest between day 3 and day 5 post-inoculation. Uninfected CTRL animals did not show signs of respiratory disease. Histopathology of the maternal lungs and placentas was evaluated to determine if IAV inoculation induced influenza-associated pathology in either organ (Fig. 1, Table S2). Lung histopathology revealed pneumonia in all cases. Of the CTRL animals with histopathology data available, none had pulmonary inflammation.

**Table 1.**
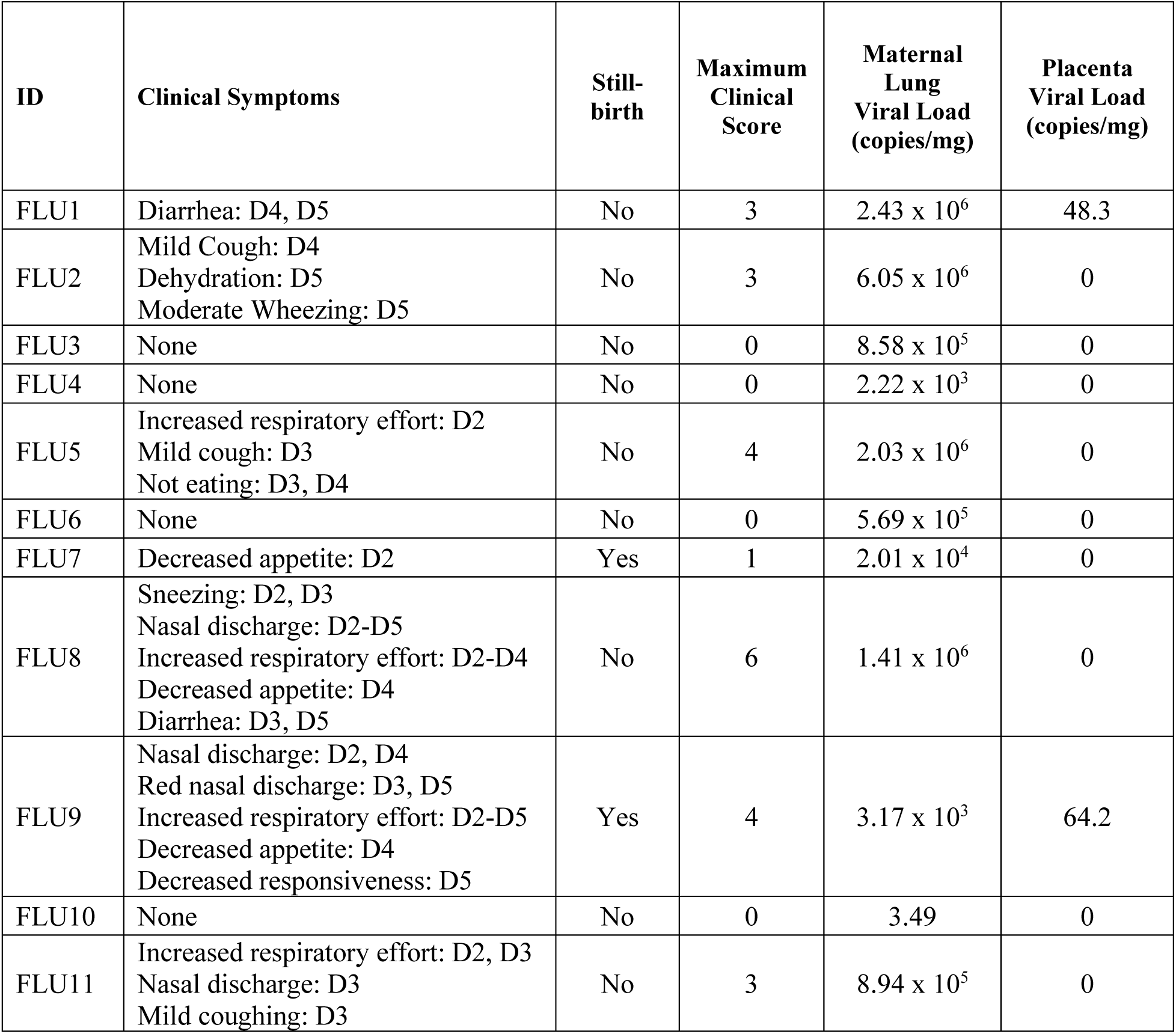
Clinical Outcomes and Maternal Lung Viral Load and Select Cytokines. Clinical symptoms were scored on a 3-point scale, with higher scores indicating worse disease and 0 indicating normal. The maximum clinical score was calculated by the sum of the highest recorded scores for each of the 5 clinical areas. Abbreviations: D, day.

**Figure 1.**
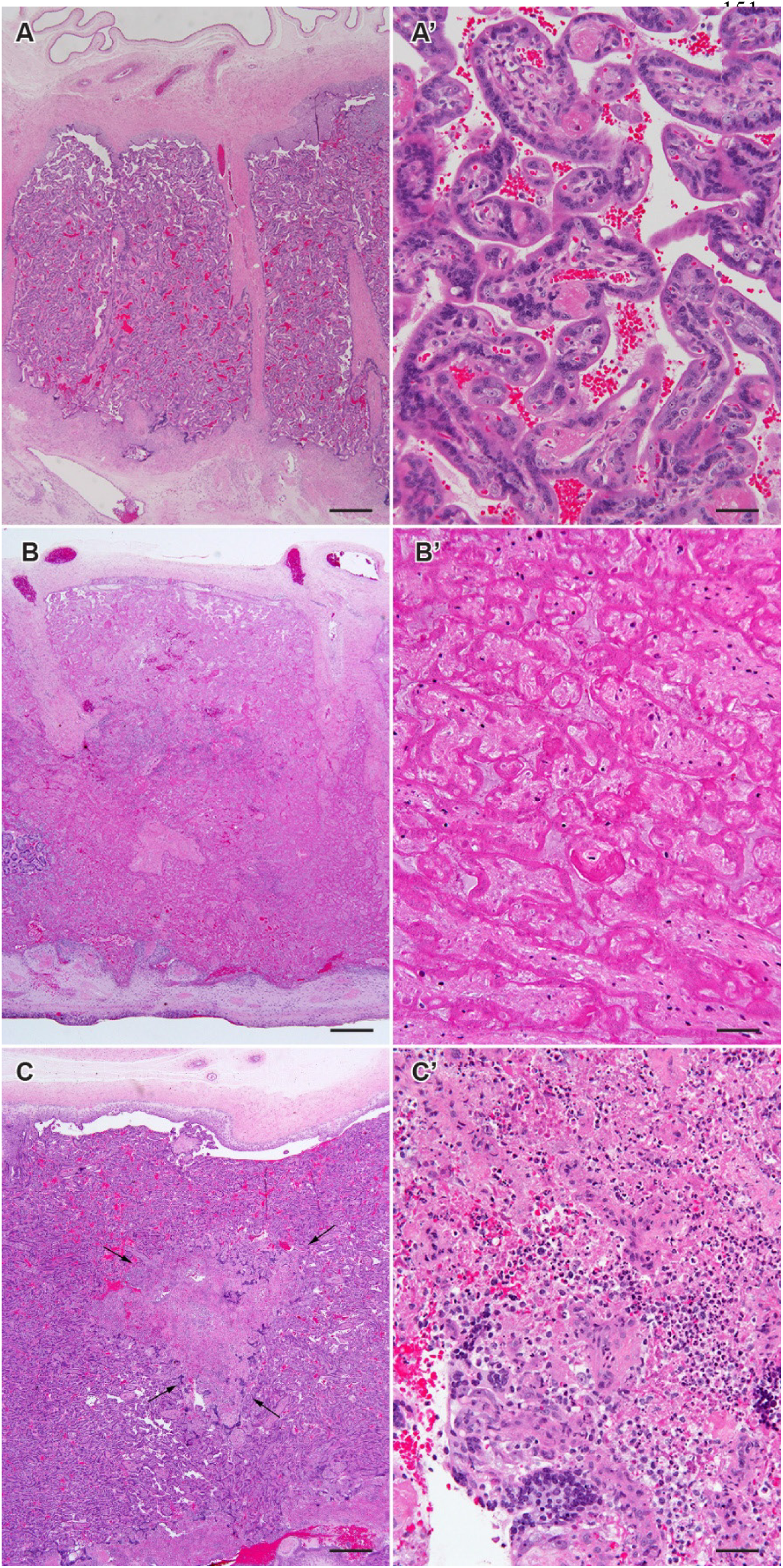
Placental Histopathology. Each pair of images shows a low-magnification image of the placental disc (fetal surface toward the top) on the left and a higher-magnification image of villous tissue on the right. **(A, A’)** Normal placental anatomy from a control animal showing no inflammation or necrosis. **(B, B’)** A focus of subacute infarction with coagulative necrosis and no active inflammation is shown from a representative animal in the FLU cohort. **(C, C’)** A focus of necrotizing villitis (highlighted by arrows) with neutrophilic inflammation and villous necrosis is shown from a representative animal in the FLU cohort.

Stillbirths were diagnosed at the study endpoint in 2 of 11 FLU animals (18%, FLU7 and FLU9), but in no CTRL animals. The gestational ages of the stillborn fetuses were 134 days (FLU 7) and 121 days (FLU 9), corresponding to ∼74% of a full-term pigtail macaque pregnancy (∼172 days gestation); the equivalent in a human pregnancy is ∼ 28 weeks gestation based on an average of 38 weeks post-conception until full-term. In nearly 6 years of data from the Washington National Primate Research Center (01/01/2019 - 12/07/2025), stillbirth occurred in 2.6% (16/608) of pregnancies viable in a comparable time window (110-140 days). The rate of stillbirths in our FLU cohort was significantly greater than that observed in the breeding colony during this gestational age time window (p=0.04).

Placental histopathology in the two stillbirth cases (FLU7, FLU9) overlapped with that observed in other IAV-infected and some CTRL animals. None of the placentas demonstrated active inflammation of the chorioamniotic membranes. FLU animals had a greater, but non-significant rate of placental infarction versus CTRL animals (FLU: 3/11, 27%; CTRL: 1/16, 6%; p=0.08). Neutrophilic (necrotizing) villitis was observed frequently in both groups (FLU: 5/11, 45%; CTRL: 4/16, 25%; Fig. 1B-C, Table S2) with no significant differences between FLU and CTRL cohorts. These findings are in line with other reports of high background rates of marginal placental infarctions and necrotizing villitis in macaque placentas.(*35-37*)

### IAV H1N1 Viral RNA Detection in Maternal Lungs, Placenta, and Fetus

RT-qPCR was performed to measure IAV viral RNA (vRNA) in the maternal lungs, placenta, and fetal organs (Table 1). At 5 dpi, all FLU animals had high levels of IAV vRNA in their lungs, ranging from 2,220 copies per milligram to more than 6 million copies per milligram; one exception was FLU10, which had an almost undetectable lung viral load (3 copies/mg). Maternal plasma was negative for IAV vRNA. In the placental chorionic villous tissue, IAV vRNA was detected in only two cases (2/11; 18%), both with a low copy number in the placenta (FLU1 and FLU9; Table 1). Of these, FLU9 additionally experienced stillbirth. Fetal tissue samples from the brain, brainstem, spinal cord, meninges, heart, lungs, liver, spleen, and thymus were negative for IAV H1N1 vRNA across the entire FLU cohort (Table S3). Vertical transmission could not be confirmed in the FLU 9 stillbirth due to severe autolysis of the tissue, rendering the RNA quality insufficient for RT-qPCR. Overall, the maternal lung was characterized by a wide distribution of IAV viral load, infrequent detection of IAV vRNA in placental tissues, and no evidence for congenital infection.

### Placental and Amniotic Fluid Cytokines

Cytokine responses in the placental disc and amniotic fluid were evaluated to determine if there was a correlation between maternal IAV infection status, viral load, or stillbirth. Chorionic villous tissue from the placenta disc of FLU animals had significantly greater concentrations of some inflammatory and antiviral cytokines than CTRL placentas, including IL-1β, IL-18, IFN-β, and GM-CSF (all p≤0.03, Fig. 2). The highest concentrations of IFN-α, IFN-β, IL-6, TNF, and GM-CSF within the FLU cohort were observed in both placentas with a low IAV viral load, of which one was a stillbirth case; however, the second stillbirth case had low cytokine concentrations. Several cytokines with low expression (IL-2, IL-10, IL-12p70, IL-13, IL-17A, and IL-23) were significantly higher in the CTRL versus FLU cohort (Fig. S1). However, in many cases, these cytokine values fell below the working range of the Luminex assay.

**Figure 2.**
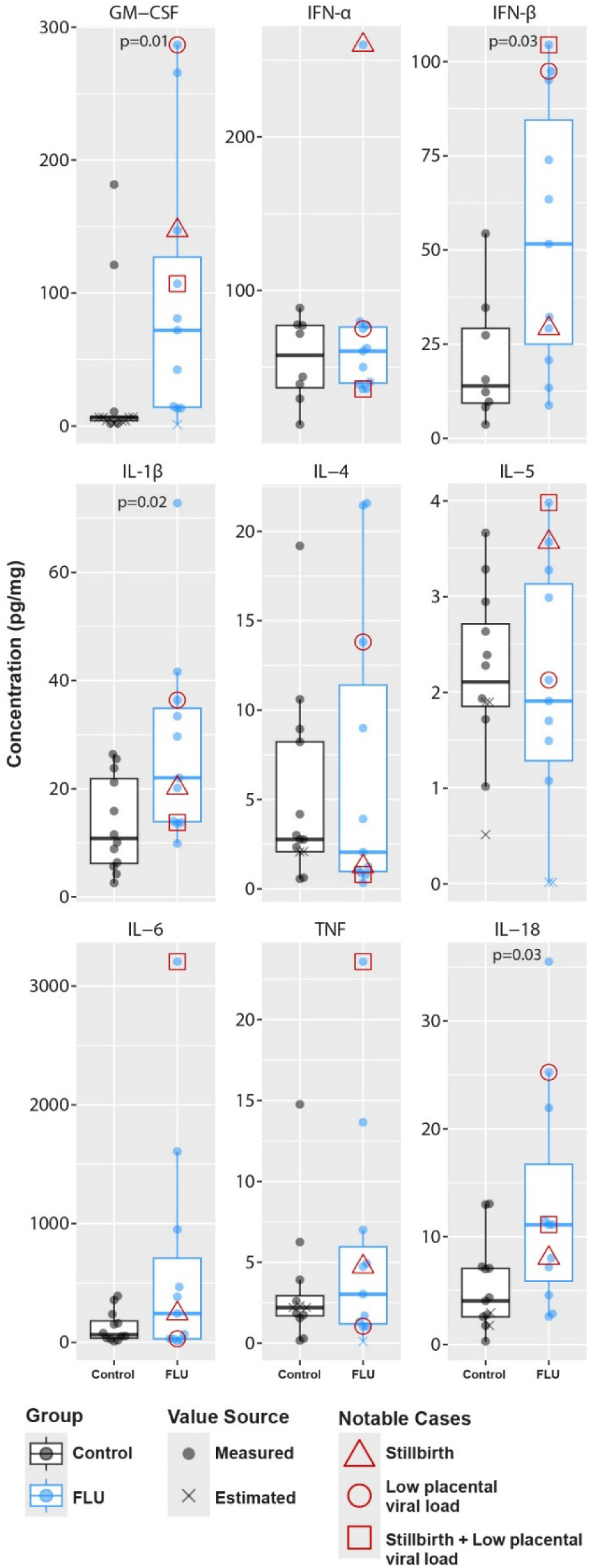
Placental Cytokine Concentrations. Each panel shows the concentration of a cytokine, quantified by a multiplex Luminex assay, in chorionic villous tissue from the FLU (blue) and CTRL (black) cohorts. Values are either measured (dots) or estimated (x’s) by extrapolation of the standard curve. Estimated values were outside the range of the standard curve and were determined by the Luminex analysis software. The thick line indicates the median, and the bounding box indicates the 1^st^ and 3^rd^ quartiles. Red triangles mark samples from animals that experienced a stillbirth. A red circle marks samples from animals that had a low placental viral load. A red square marks samples from animals that experienced a stillbirth and had low placental viral load. P-values are shown using a Wilcoxon rank sum test comparing FLU versus CTRL cohorts.

In contrast to chorionic villous tissue, several cytokines in the amniotic fluid were significantly higher in the CTRL versus FLU cohort (GM-CSF, IFN-β, IFN-γ, IL-1β, IL-4, IL-5, IL-13, IL-17A, and IL-23; all p<0.05). In most cases, cytokine concentrations in both cohorts were very low (Fig. S2). These results indicate that perturbations in cytokines associated with maternal IAV infection were compartment-specific, and elevated cytokines in the chorionic villous tissue were not transported into the amniotic fluid compartment.

### Placental Immune Cell Frequencies

To determine whether maternal IAV H1N1 infection was associated with changes in the innate and adaptive immune cell composition of the placental disc, we performed flow cytometry-based immunophenotyping. The frequency of classical monocytes among live leukocytes was significantly decreased in the placental disc of the FLU versus CTRL cohort (p=0.004, Fig. 3). There was no relationship between stillbirth and the frequency of immune cell populations in the placenta. To determine the impact of gestational age matching, we performed a more conservative analysis that included only CTRL animals with gestational age at endpoint within +/- 1 week of the average gestational age of the FLU cohort (Fig. S3). When restricting the analysis to gestational age-matched controls, the distributions of CD4+ T cell, CD8+ NK cell, CD8+ T cell, and classical monocyte frequencies were significantly lower in FLU versus CTRL cohort (p=0.02-0.04, Fig. S3).

**Figure 3.**
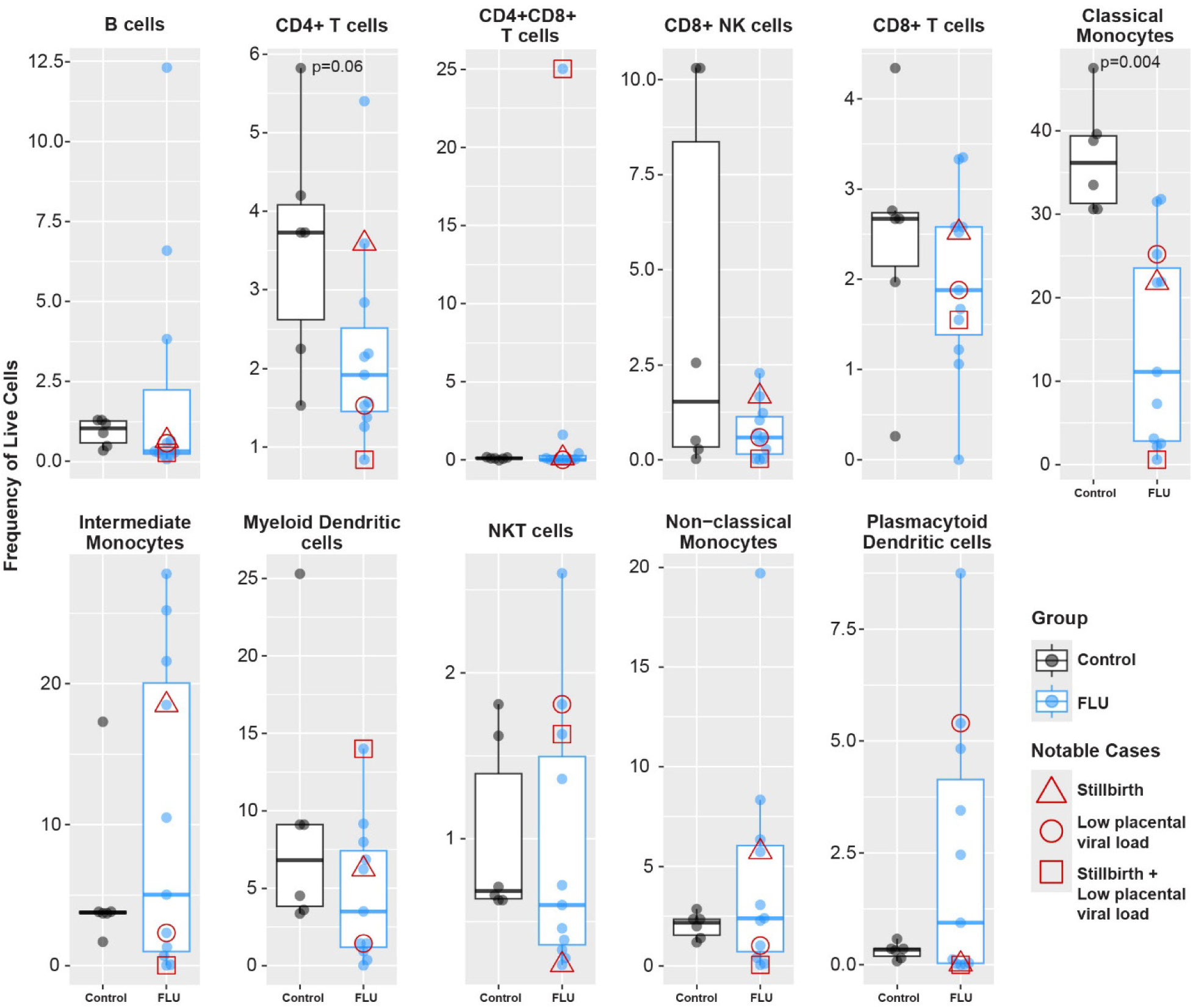
Placental Immunophenotyping. Each panel shows the frequency of the defined cell population as a percentage of live cells, in chorionic villous tissue of the FLU (blue) and CTRL (black) cohorts. The thick line indicates the median, and the bounding box indicates the 1^st^ and 3^rd^ quartiles. A red triangle marks samples from animals that experienced stillbirth. A red circle marks samples from animals that had a low placental viral load. A red square marks samples from animals that experienced a stillbirth and had low placental viral load. P-values are shown using a Wilcoxon rank sum test comparing FLU versus CTRL cohorts. Significance is considered for p-values < 0.05 but is shown if 0.05 ≤ p ≤ 0.1 due to the modest sample size. Abbreviations: NKT, natural killer T cells

### Placental Transcriptional Profile

To determine if a maternal IAV infection altered the transcriptional signature in the placental disc, we performed total RNA-Seq on chorionic villous tissues and validated our results by direct gene counting using a Nanostring nCounter assay. A multidimensional scaling plot was created to visualize the normalized RNA-Seq gene counts and demonstrated a clear division between the two cohorts (CTRL, blue dots) and FLU (red symbols; Fig. 4A). In the placental chorionic villous tissue, 185 genes were upregulated, and 178 genes were downregulated relative to the gene expression of the CTRL group (≥ or ≤ 1.5-fold, p<0.05; Fig. 4B). Inspection of the upregulated genes revealed a striking upregulation of interferon (IFN)-stimulated genes (ISGs) including *IFIT3, MX1*, and *OAS2* in placentas from FLU animals relative to CTRL placentas. A Gene Set Enrichment Analysis (GSEA) revealed multiple significant, positively enriched pathways related to the type I IFN response and the negative regulation of viral processes such as genome replication and viral entry in FLU versus CTRL animals (Fig. 4C). Digital gene counting using a 770-plex Nanostring nCounter panel supported these results with a similar striking upregulation of ISGs (Fig. S4), and strong correlation with total RNA-Seq gene counts in both the FLU and CTRL groups (Fig. 4D; r=0.8, p<2.2×10^-16^ for both).

**Figure 4.**
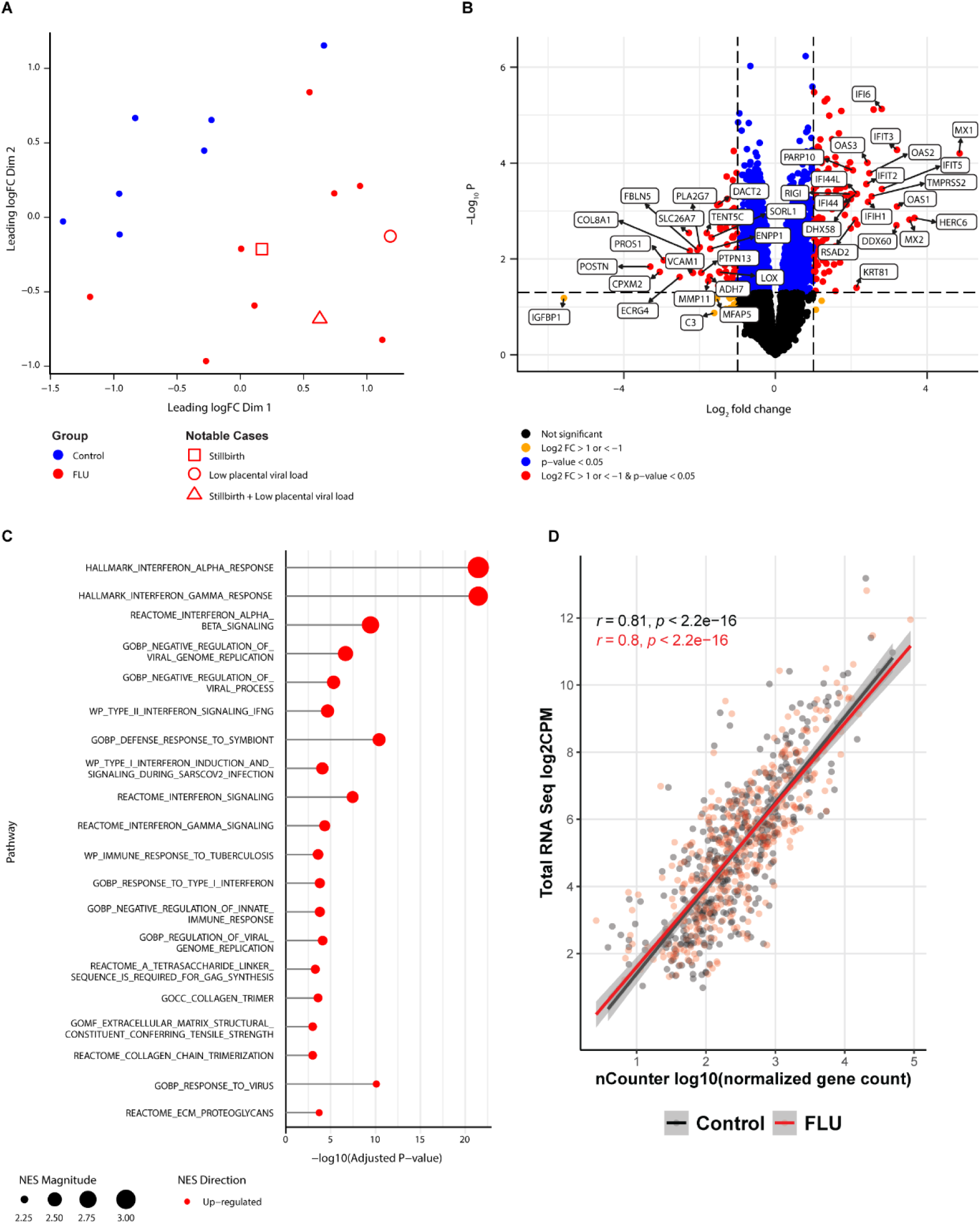
Placenta Total RNA-Seq Analysis. **(A)** Multidimensional scaling (MDS) of normalized log-transformed gene counts using Euclidean distance depicts sample-to-sample similarity; each point represents a placenta and is colored by group (FLU: red; CTRL: blue). Open red symbols denote FLU samples with low placental IAV vRNA (circle), a stillbirth with undetectable vRNA (square), or a stillbirth with low vRNA (triangle). **(B)** The volcano plot shows log2 fold change versus –log10(p-value) for FLU versus CTRL. Red dots mark significant genes with |log2FC| ≥1; blue mark genes with |log2FC| <1 but p<0.05; gold indicate non-significant genes with |log2FC| ≥1; black indicate non-significant genes with |log2FC| <1. **(C)** Gene Set Enrichment Analysis of significant GO terms (p<0.05) displays the normalized enrichment score (NES) for each category; FDR values reflect correction for multiple testing **(D)** The dot plot shows correlations between placental log2-transformed RNA-Seq counts per million and log10-transformed Nanostring nCounter counts. Each point is the mean expression of a gene present in both datasets from FLU (red) or CTRL (black) samples. Pearson r and p-values are shown, with the best-fit line and standard error.

Next, we compared bulk transcriptomic changes in the placenta between the FLU and CTRL cohorts to identify pathways differentially expressed by IAV. Hierarchical clustering yielded two distinct clusters of differentially expressed genes: cluster 1 (downregulated genes, FLU vs. CTRL; Fig. 5A) and cluster 2 (upregulated genes, FLU vs. CTRL; Fig. 5A). Overrepresentation Analysis (ORA) was performed to identify Gene Ontology (GO) terms enriched in both clusters. No significant GO terms were identified in cluster 1. However, inspection of genes in cluster 1 revealed downregulation in the FLU group of multiple genes involved in ubiquitination, proteasome function, unfolded protein response, metabolism, and cell cycle/DNA repair (Fig. S6). To determine whether this profile might reflect the induction of a viral-associated integrated stress response (ISR), we analyzed the expression of two key pathway genes: *EIF2AK2* (ISR activator) and *ATF4* (ISR effector). Of these genes, *EIF2AK2* expression was significantly upregulated in the placenta [log2 fold change=1.24, p<0.05; Fig. S6]. In cluster 2, significantly enriched GO terms in the FLU cohort were related to type I and II interferon signaling, viral genome regulation, and antiviral defense processes that featured ISG genes (Fig. S5, S7). A notable outlier was FLU11, which had a transcriptional profile more like that of CTRL animals. Placental transcriptomics of FLU3 and FLU4 were intermediate between the FLU and CTRL cohorts. Collectively, transcriptional analysis of the placental chorionic villous tissue from FLU animals revealed a striking upregulation of antiviral innate immune genes and downregulation of critical cellular functions (cell cycle, metabolism, ubiquitination) consistent with an antiviral and ISR response, regardless of the severity of disease in the maternal lung.

**Figure 5.**
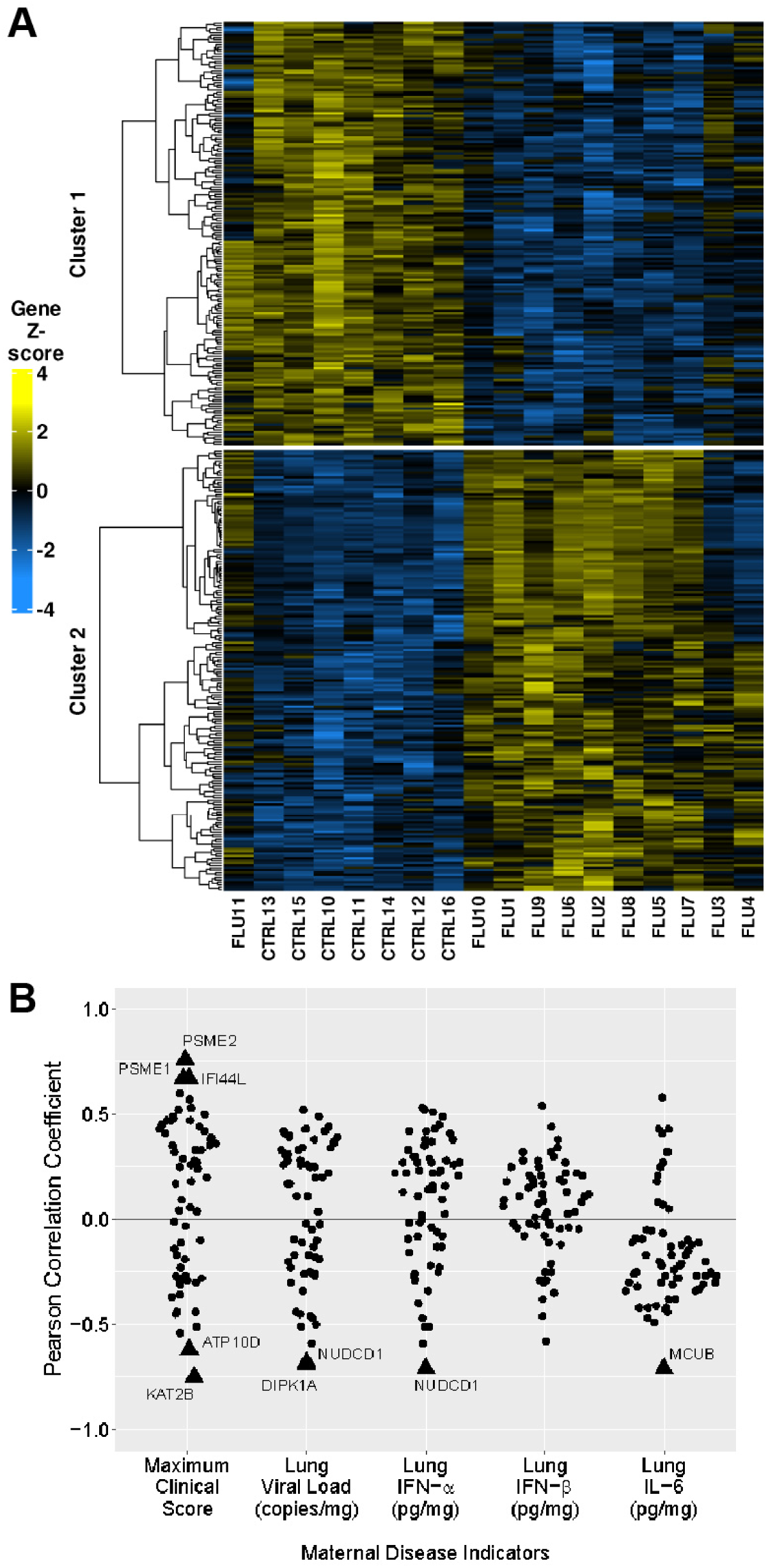
Placental Total RNA-Seq Clustering and Correlation Analysis. **(A)** Heatmap of normalized gene counts from placental total RNA-seq comparing FLU-exposed and CTRL cohorts. Each column represents an individual placental sample, and each row represents a single gene. Yellow-to-blue color intensity reflects the gene-level z-score, calculated using the mean expression value across all animals for each gene. Unsupervised hierarchical clustering was performed on the top 500 differentially expressed genes identified in FLU placentas, using Euclidean distance and Ward’s linkage method. **(B)** Pearson correlation analysis between a select panel of 64 placental genes (listed in Fig. S6-S7) and key metrics of maternal IAV disease severity: maximum clinical score, lung viral load, lung IFN-α concentration, lung IFN-β concentration, and lung IL-6 concentration. Cytokine concentrations and viral load were log10-transformed before analysis. The Pearson correlation coefficient (R) is plotted for each gene-metric pair. Significant correlations (p<0.5) are shown as triangles, and non-significant correlations (p ≥0.5) are shown as dots. Genes with statistically significant correlations are labeled.

### Placental Correlations with Maternal Lung Disease Indicators

We hypothesized that placental gene expression might correlate with either clinical score, lung viral load, or an immunologic correlate of IAV-associated lung disease, such as an inflammatory cytokine (IL-6), or the antiviral type I IFN response (lung IFN-α, IFN-β). A panel of 64 key genes derived from the hierarchical clustering model was selected as key genes for the correlation analysis (listed in Fig. S6, S7). Few correlations between these outcome metrics emerged (Fig. 5B). Evaluating correlations between maternal disease indicators and placental cytokines or immune cell frequencies similarly yielded almost no significant correlations (Fig. S8-S9). Overall, this data suggested a weak correlation between IAV-associated maternal lung disease indicators and placental gene expression or cytokine concentrations.

### Fetal Immune Cell Frequencies in Lung, Lymph Node, and Blood

Next, we examined fetal lung, lymph nodes, and blood compartments using flow cytometry-based immunophenotyping. In these fetal organs, there was a coordinated perturbation of innate immune cells and T cells in the FLU versus CTRL cohorts. In the fetal lung, the frequencies of intermediate monocytes, non-classical monocytes, plasmacytoid dendritic cells, and CD4+ T cells were significantly higher in the FLU cohort compared to the CTRL cohort (p≤0.04, Fig. 6). A similar pattern was evident in the fetal lymph nodes, with a significantly higher frequency of intermediate monocytes (p=0.02). Additional immune populations in the FLU versus CTRL fetal lymph node had a higher, but non-significant distribution of NKT cells, non-classical monocytes, and plasmacytoid dendritic cells (all p=0.06, Fig. 6). In fetal whole blood, there was a significantly higher frequency of NKT cells in the FLU cohort (p=0.03, Fig. 6). After restricting these analyses to CTRL animals with data obtained +/- 1 week of the average FLU cohort, the significant findings of higher frequencies of intermediate and non-classical monocytes in the fetal lungs, and a higher frequency of NKT cells in whole blood remained (p=0.004 - 0.01; Fig. S13). Overall, these differences suggest that maternal IAV H1N1 infection perturbs populations of fetal innate and adaptive immune cells in organs, lymphoid compartments, and blood, even in the absence of vertical transmission.

**Figure 6.**
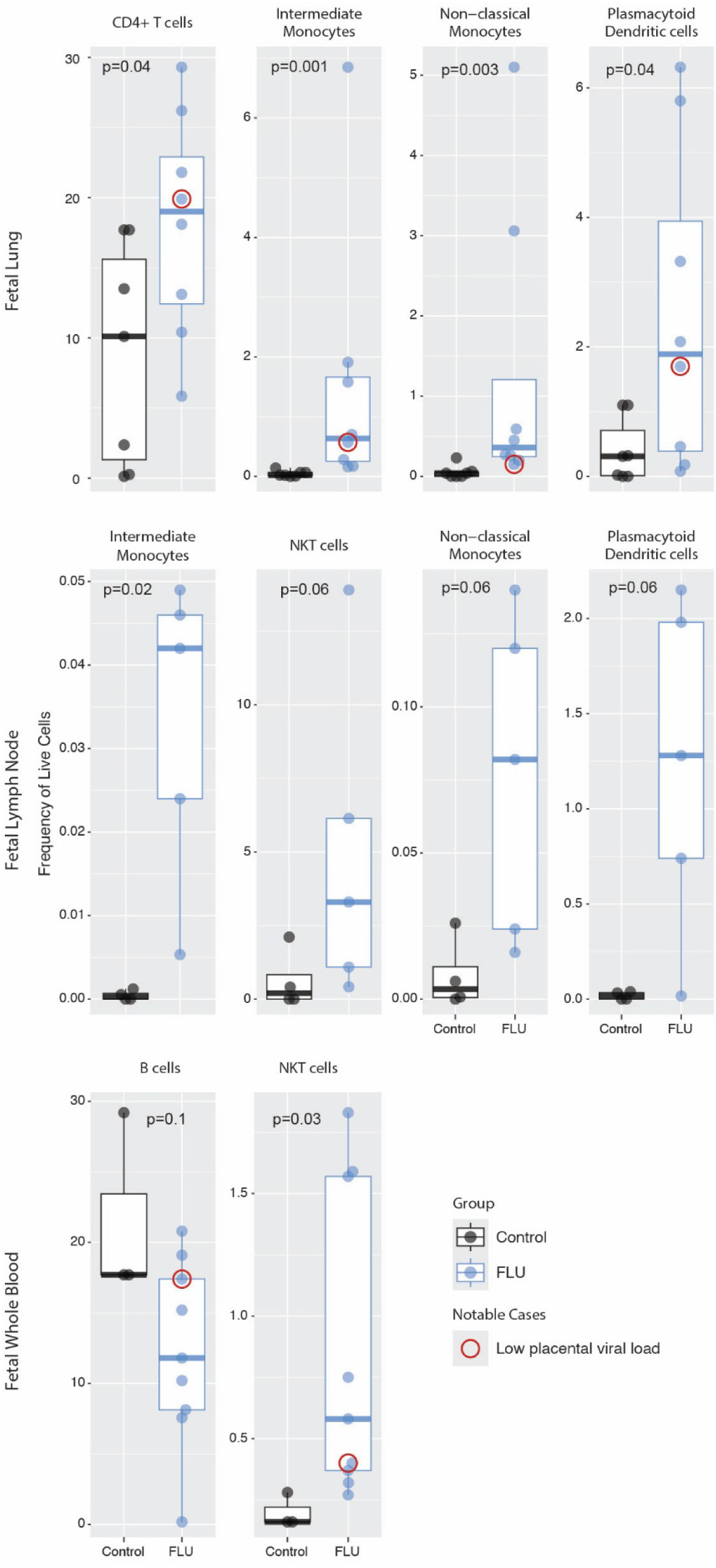
Fetal Immunophenotyping. Each panel shows the frequency of the defined cell population as a percentage of live cells, in fetal lung (top row), fetal lymph node (middle row), and fetal whole blood (bottom row) of the FLU (blue) and CTRL (black) cohorts. The thick line indicates the median, and the bounding box indicates the 1^st^ and 3^rd^ quartiles. A red circle marks samples from animals that had a low placental viral load. Fewer number of observations in a group reflects inability to obtain sufficient sample or cells to perform the assay. Immunophenotyping was not done on fetal samples from stillbirth cases. The p-values were generated using a Wilcoxon rank-sum test and are indicated at the top of each panel. Significance is considered for p-values < 0.05 but is shown if 0.05 ≤ p ≤ 0.1 due to the modest sample size. Additional results are in Fig. S10-S12. Abbreviations: NKT, natural killer T cells

### Fetal Correlations with Maternal Lung Disease Indicators

Next, we were interested in exploring correlations between maternal lung disease indicators with fetal immune cell populations. In contrast to the few correlations identified in the placenta, there were multiple significant correlations between IAV-associated maternal lung disease and fetal immune cell populations that remained consistent across diverse organ and fluid compartments (lungs, lymph nodes, and whole blood, Fig. 7). A common pattern emerged of significant, positive correlations between metrics of maternal disease severity and higher frequencies of fetal T cell populations (CD4+, CD8+, CD4+CD8+ T cells) or plasmacytoid dendritic cells, which are major producers of type I IFNs (all, p<0.05). Interestingly, maternal disease indicators were negatively correlated in some cases with fetal monocyte populations (classical, non-classical, intermediate). Collectively, these data suggest that maternal IAV disease is associated with perturbations in immune cells across multiple fetal tissue and fluid compartments.

**Figure 7.**
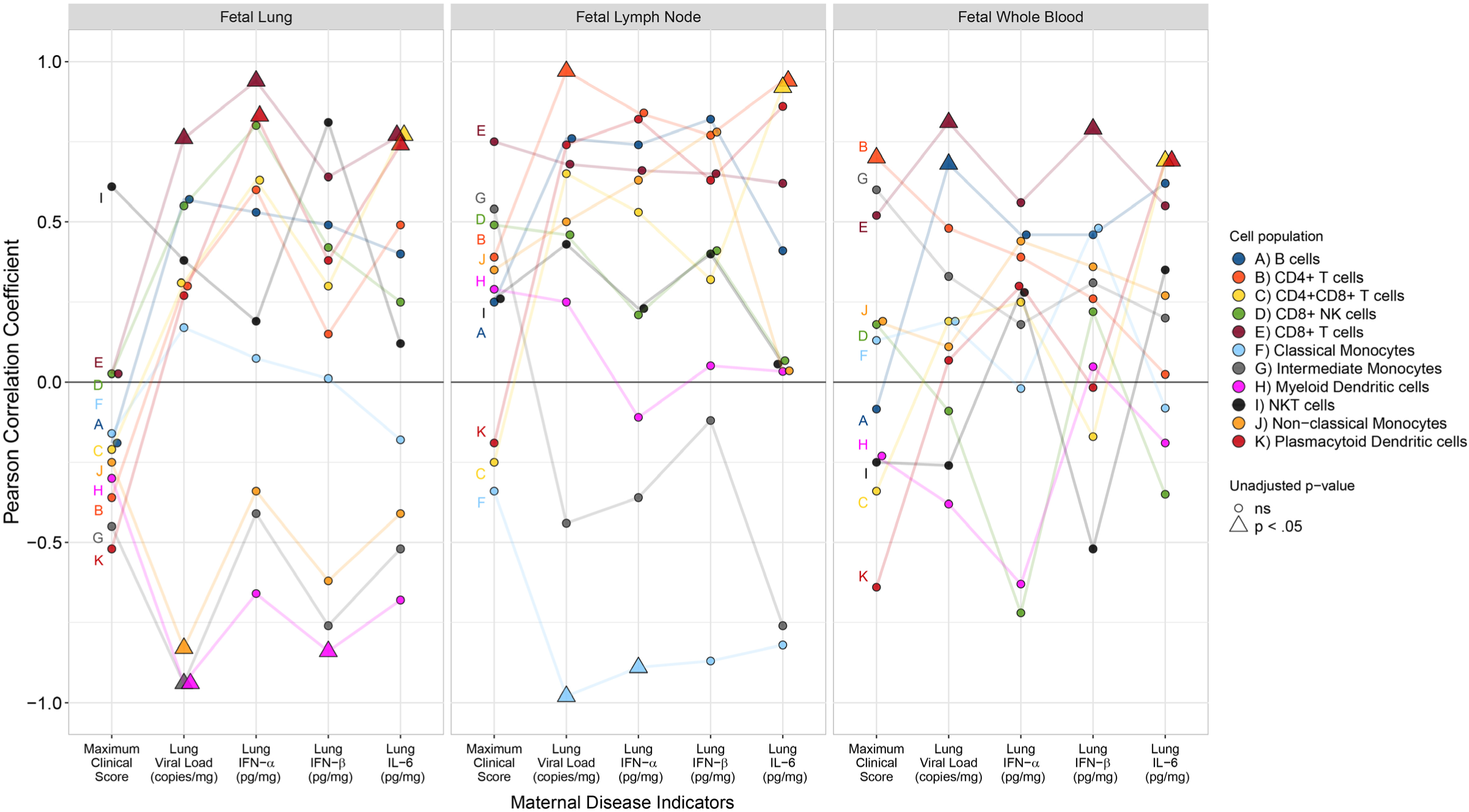
Pearson Correlation Coefficients Between Maternal Lung Disease Indicators and Fetal Immune Cell Populations. Pearson correlation analysis between maternal disease indicators (maximum clinical score, lung viral load, lung IFN-α/IFN-β/IL-6 concentration) and fetal immune cell populations in the fetal lungs, lymph nodes, and whole blood. The Pearson correlation coefficient is shown on the y-axis, and the maternal disease indicator is shown on the x-axis. Each immune cell population is indicated by a color and a letter to aid in distinguishing them. A line connects each Pearson correlation coefficient from the same immune cell population, allowing tracking of similarities in positive and negative coefficients. Triangles show significant unadjusted p-values.

## DISCUSSION

The placenta is a critical immunological gatekeeper during pregnancy, balancing its essential roles in nutrient and oxygen exchange with the need to protect the fetus from invading pathogens. Our data suggest that placental IAV exposure induces a cascade of immune defenses, including a robust placental type I interferon response (IFN-β) and activation of inflammasome pathways (IL-1β, IL-18). These responses are characteristic of a rapid antiviral defense state, in which the placenta prioritizes pathogen restriction over immune quiescence to protect and nourish the fetus. Importantly, this placental antiviral response occurred independently of maternal lung disease severity, supporting the concept that the placenta transitions into an autonomous defensive state in response to maternal infection, effectively shielding the fetus from infection at the potential cost of localized immune activation, placental injury, and stillbirth.

To our knowledge, this study provides the first demonstration that maternal IAV infection can broadly perturb the fetal immune landscape in a manner linked to maternal disease severity. As vertical transmission was not detected in our experiments, the data suggest that immunologic, rather than virologic, signals are responsible for the profound changes in the fetal immune landscape observed after in utero exposure to maternal IAV infection. Notably, fetuses in the FLU cohort exhibited significantly higher frequencies of multiple T-cell populations and reduced frequencies of monocytes compared with CTRLs, with these changes evident across multiple organs and fluid compartments and correlated with key metrics of maternal IAV disease. Although the mechanisms by which host–pathogen interactions shape fetal immune development and long-term health remain incompletely understood, maternal disease severity represents a potentially modifiable determinant of these outcomes.

Altered fetal immunity has previously been described following in utero exposure to maternal human immunodeficiency virus (HIV) infection, even in the absence of maternal–fetal viral transmission. Perinatally HIV-exposed but uninfected infants have been shown to have a lower frequency of naïve CD4⁺ and CD8⁺ T cells, among other abnormalities in immune activation, compared to perinatally HIV-unexposed infants.(*38-41*) Similar findings of a perturbed fetal immune landscape have been reported in cases of maternal severe acute respiratory syndrome coronavirus 2 (SARS-CoV-2) infection. Perturbed monocyte cell populations with increased NK cell and regulatory T cell frequencies were reported in uninfected neonates exposed to maternal SARS-CoV-2.(*42, 43*) These changes in fetal immunity may represent an intermediate biological state that accompanies pathways leading to abnormal neurodevelopment, as described in many cohorts of adults exposed to maternal influenza and other infections during fetal life.(*13, 14, 19, 20, 44, 45*)

Stillbirth occurred in two animals in the FLU cohort, an outcome that was unexpected based on the CTRL cohort and historical colony experience. The placental antiviral program triggered by IAV H1N1 is known to promote inflammation and tissue injury during acute viral infections, and has been directly implicated in the pathogenesis of stillbirth in mouse models of congenital Zika virus infection.(*46-49*) Whether a viral-associated ISR may have also contributed to stillbirth through translational repression and maladaptive stress signaling is unknown. We are cautious to causally attribute these stillbirths solely to the placental type I IFN or ISR response for several reasons. A spectrum of placental pathology was observed in FLU animals, with similar lesions (i.e., infarction, necrotizing villitis) also present in several CTRL placentas from clinically healthy animals. Background placental pathology in macaques is known to be relatively high, with prior studies documenting marginal infarctions and villitis as common findings even in uncomplicated pregnancies.(*35-37*) Moreover, unlike human stillbirth evaluations, which routinely involve extensive histopathologic sampling across the entire placental disc, the placental tissue in this study was allocated to multiple downstream immunologic and transcriptomic analyses, thereby limiting a comprehensive histopathologic examination. Together, these factors limit our ability to assign a specific pathological mechanism to the two stillbirths, underscoring the need for cautious interpretation while highlighting a biologically plausible link between antiviral placental immune activation and adverse perinatal outcomes.

The primary strength of our study lies in the translational nature of the NHP model, which closely recapitulates human pregnancy, enabling modeling of an IAV infection in the context of pregnancy. NHPs are highly similar to humans in their genetics, anatomy, physiology, immune responses, and pregnancies.(*27*) Macaques have a long gestational period and a placental architecture similar to that of humans. Another strength of the pregnant NHP model of IAV infection is the ability to study antiviral immune responses at a defined time after infection onset, which is not possible during human pregnancy. Further, this study was deliberately conducted in the third trimester of pregnancy, a time when pregnant women experience the most significant maternal morbidity and mortality.(*50, 51*) NHP studies excel at identifying integrated physiologic outcomes that cannot be replicated in other models, providing an essential translational context for human pregnancy. Defining molecular mechanisms in NHP models is constrained by biology and ethics, in contrast to *in vitro* or mice models, which offer significantly greater experimental manipulation. These features of the pregnant NHP model of IAV infection demonstrate the translational relevance of this model and the marked impact of our findings, while also providing possible mechanistic insights for future studies.

This study has several limitations, primarily related to ethical constraints involved with NHP studies and the modest sample size. We did not study endpoints earlier or later than five days post-infection, preventing assessment of the trajectory of the antiviral response in the maternal lungs and placenta. We used multiple routes of viral inoculation to ensure productive disease; experiments using lower inocula delivered via aerosol would more closely recapitulate IAV transmission in humans. Examining multiple endpoints was not feasible with our limited sample size, but future studies could investigate alterations in fetal immune regulation after maternal IAV disease resolution and the potential for development of an integrated stress response in the placenta. While the stillbirths are notable cases, the study’s conclusions are drawn from patterns across the entire infected cohort. Our CTRL cohort consisted of historical uninfected controls used in prior studies. While housing conditions, sample collection, and experimental protocol remained consistent across all animals, the use of historical controls provides challenges in accurately controlling confounding variables.

Together, these findings position the placenta as an active and autonomous organ that mounts a robust type I IFN and inflammasome-mediated defense against IAV. While this response appears effective at preventing vertical transmission, it is accompanied by localized immune activation and placental injury that may contribute to adverse pregnancy outcomes. In contrast, fetal immunity is sensitive to maternal disease severity, with perturbations in fetal T cells and monocytes across multiple fetal compartments occurring in a coordinated response. Despite public health recommendations for pregnant women to receive the seasonal IAV vaccine (51), only half of pregnant women in the U.S. typically receive it each year.(*52*) Seasonal IAV vaccination is known to reduce maternal morbidity, mortality, and stillbirth.(53–59) Whether mitigation of maternal IAV disease through vaccination preserves the normal trajectory of fetal immune development is an important question.

## MATERIALS AND METHODS

### Study Design

Two groups of female pigtail macaques *(Macaca nemestrina*) were inoculated using a combination of intranasal, intratracheal, intraocular, and intraoral routes with 10^7.4^ PFU of IAV [A/California/07/2009 (H1N1)], reflecting a dosing method previously used in a study of cynomolgus macaques (*Macaca fascicularis*), which resulted in a productive infection.(*53*) Experimental groups consisted of IAV H1N1-infected pregnant (N=11) and uninfected pregnant controls (N=18; Table S1). IAV inoculation occurred in the early third trimester to model the time in pregnancy when IAV H1N1 infections are thought to produce the greatest maternal lung disease and pregnancy complications.(*11, 12*) Uninfected controls in this study consisted of pregnant adult pigtail macaques used in prior published studies from two cohorts: 1) control studies catheterizing the amniotic cavity and inoculating saline into the choriodecidual space and/or amniotic fluid; 2) control studies inoculating media subcutaneously to mimic procedures in a Zika virus experimental cohort.(*54, 55*) Ethical constraints limit the availability of large, dedicated control cohorts for each experimental study. Thus, historical controls were used with consistent husbandry conditions, necropsy and processing protocols, and experimental protocols to minimize potential confounding variables. Animals in the WaNPRC were not exposed to seasonal IAV infections and thus do not have pre-existing immunity to IAV.

### Ethics Statement and Animal Care

All animal experiments used in this study were approved by the University of Washington Institutional Animal Care and Use Committee (protocol# 4165-03, last approved 07/22/2025), and in compliance with the U.S. Department of Health and Human Services Guide for the Care and Use of Laboratory Animals and Animal Welfare Act. All animal experiments were performed at the Washington National Primate Research Center (WaNPRC).

Macaques inoculated with IAV were observed daily for clinical symptoms of influenza disease, which were scored on a 3-point scale to reflect responsiveness of the animal, nasal or ocular discharge, cough, respiratory effort, food consumption, and fecal consistency; higher scores reflected disease, with a 0 in each category indicating a normal assessment. We chose a predefined study endpoint of 5 days post-inoculation (dpi) to capture immune responses near the expected peak of disease.^35,36^ A Cesarean section was performed at the endpoint, and placentas were extracted to sample chorionic villous tissue. The dam was euthanized using a barbiturate overdose while under anesthesia. The fetus was euthanized using a barbiturate overdose, followed by exsanguination. These methods are approved by the 2007 American Veterinary Medical Association Guidelines on Euthanasia. Necropsies were performed on both the dam and the fetus to obtain samples for transcriptomics, immune assays, and histopathology. Random sections of the placental membrane, chorionic disc, and umbilical cord were evaluated for pathology. In one stillbirth case, FLU 9, the fetus was 1-2 days post-mortem at the time of Cesarean section. Pathologists reported that severe autolysis of the tissue had occurred. Samples were not collected from the fetus because the tissue quality would likely be too low to perform meaningful assays.

### IAV H1N1 Virus Cultivation and Quantification

We used an IAV strain [A/California/07/2009 H1N1 influenza virus (CA09)] that was obtained from the World Reference Center for Emerging Viruses and Arboviruses at the University of Texas Medical Branch at Galveston. Before culturing the virus, Madin-Darby canine kidney (MDCK) cells (ATCC, Manassas, Virginia, US) were cultured in Dulbecco’s modified Eagle’s medium (DMEM) supplemented with 10% fetal bovine serum (FBS), 25 mM HEPES, 100 U/mL penicillin, and 100 μg/mL streptomycin. Cells were incubated at 37°C and 5% CO_2_ and confirmed to be free of mycoplasma with the MycoAlert™ mycoplasma detection kit (Lonza, LT07-318) before use for virus cultivation. Sub-confluent monolayers of MDCK cells in T150 flasks were washed with PBS and infected at a multiplicity of infection (MOI) of 0.5 (5 mL serum-free MEM) at 37°C and 5% CO_2_. After 1 hour, the media was removed, cells were washed with 1X PBS, and the media was replaced. The supernatant was harvested after 36 hours and centrifuged at room temperature to remove cellular debris (1800 rpm for 5 minutes, followed by 3000 rpm for 10 minutes). Virus stock was frozen at −80°C. The viral titer of each stock was quantified using a TCID50 assay with hemagglutination readout and converted to PFU/mL.(*56*) To transform TCID50/mL into PFU/ml, TCID50/mL was multiplied by 0.7. Reagents were obtained from Thermo Fisher Scientific unless otherwise stated (Waltham, MA).

### RNA Isolation and Reverse Transcription Quantitative Polymerase Chain Reaction (RT-qPCR)

Tissue samples were kept in RNAlater™ Stabilization Solution (#15596018, Thermo Fisher Scientific) prior to homogenization in Precellys homogenizing tubes (#1011152, Cayman Chemical, Ann Arbor, MI, USA) with the Precellys Evolution machine (Bertin Technologies, Montigny-le-Bretonneux, France). RNA was extracted from homogenates using the RNeasy Mini Kit (Qiagen, #74106) and treated with DNase (Qiagen, #79256). RNA was quantified using the NanoDrop™ 2000 spectrophotometer (Thermo Scientific, #ND-2000) and the Qubit 4 fluorometer (Invitrogen, #Q33238). RNA quality was evaluated by measuring the RNA integrity number (RIN) with the 4200 Tapestation System (Agilent, #G2991BA).

To quantify IAV vRNA levels from different sample origins, we used reverse transcription quantitative polymerase chain reaction (RT-qPCR) and a WHO-approved primer/probe set.(*57*) cDNA was generated from RNA isolates using an iScript™ Select cDNA synthesis kit (Bio-Rad, #1708897) using H1N1 HA-specific forward primer following manufacturer’s instructions. qPCR was done with H1N1 HA-specific primers and probes (Forward: GGGTAGCCCCATTGCAT; Reverse: AGAGTGATTCACACTCTGGATTTC; Probe: 5’-[FAM] TGGGTAAATGTAACATTGCTGGCTGG [TAMRA]-3’) using the Taqman Fast Advanced Master Mix (#444964, Thermo Fisher Scientific) and a QuantStudio 3 machine (#A28132, Thermo Fisher Scientific) following manufacturer instructions. Samples were run in triplicate.

An external plasmid standard curve was used to determine viral RNA quantities with the lower limit of quantification being 3 copies per reaction. A negative control with no template added was used to assess the possibility of contamination. Amplification was defined as fluorescent activity exceeding a threshold of 0.1. Negative results were defined as any sample that did not amplify or had a Ct value greater than 38 cycles. Positive results were defined as any amplified sample with a Ct value less than 38 cycles for all three replicates. Samples with low positive results (Ct value>35) had RNA re-extracted from a new sample aliquot and RT-qPCR was run two more times in triplicate to verify results, totaling 9 replicates. Copies of viral RNA per milligram of tissue were calculated based on tissue weight, RNA concentration, RNA volume, and volume of cDNA sample input.

### Cytokine Quantification

ProcartaPlex™ NHP multiplex kits were used to quantify cytokines from lysates or fluids (Thermo Fisher Scientific). Tissue lysis and homogenization were performed using an Invitrogen™ ProcartaPlex™ Cell Lysis Buffer (#EPX-99999-000, Thermo Fisher Scientific) with the addition of a protease inhibitor cocktail (Fisher Scientific, #PI78430) using the Precellys Evolution system, as previously described. Type I Interferons [IFN; IFN-alpha (IFN-α) and IFN-beta (IFN-β)] were quantified using the Cynomolgus IFN-beta and Cynomolgus/Rhesus IFN-alpha ELISA kits (#46415-1 and #46100-1, respectively; PBL Biosciences, Menlo Park, CA, USA). All protocols followed the manufacturer’s guidelines. The Luminex analysis software estimated cytokine concentrations that fell outside the range of the standard curve.

### Immunophenotyping

Immunophenotyping by flow cytometry was performed to determine the frequencies of immune cells in the placenta and fetal lung, mesenteric fetal lymph node, and fetal blood. Tissues from the placental disc and fetal organs were dissociated into single cell suspensions and assessed for viability using Acridine Orange/Propidium Iodide (Nexcelom Bioscience LLC). Then, samples were stained with a panel of antibodies in Brilliant Stain Buffer (BD Biosciences) for 20 minutes to identify immune cells as previously described.(*58*) Antibodies used included BD Biosciences: CD3 (SP34-2)-BV650, CD8 (RPA-T8)-BUV395, CD11b (ICRF44)-APC-Cy7, CD11c (SHCL-3)-PE, CD45 (D058-1283)-PE-CF594, CD86 (IT2.2)-PE-Cy5, TCRγδ (B1)-FITC; BioLegend: CD4 (OKT4)-BV605, CD14 (M5E2)-BV785, CD16 (3G8)-Alexa Flour 700, CD20 (2H7)-BV570, HLA-DR (L243)-BV711; Invitrogen: CD123 (6H6)-eFlour 450, LIVE/DEAD Fixable Aqua for 405nm excitation; Beckman Coulter: CD159a (Z199)-PC7; and Miltenyi: CD66 (TET2)-APC. The staining protocol proceeded in three steps: LIVE/DEAD Fixable Aqua, then all antibodies other than CD11c and CD123, and finally CD11c and CD123. After staining, samples were fixed with 1% paraformaldehyde and analyzed on a Symphony A3 flow cytometer (BD Biosciences) using FACS Diva 8 software (BD Biosciences). Samples were analyzed using FlowJo 10.9.0 (FlowJo, LLC). All events were initially gated on Forward Scatter (FSC) singlets, CD45+ leukocytes, live cells, and then on FSC-A and Side Scatter (SSC)-A profiles. Populations were defined using canonical markers as described in Table S4. Representative gating strategies are shown for granulocytes (Fig. S14), lymphoid cells (Fig. S15), and monocytes (Fig. S16). Populations were excluded from analysis if they did not meet the minimum threshold of >100 cells in their parent gate.

### Total RNA-Seq Library Preparation and Analysis

Total RNA sequencing was performed on chorionic villous tissue from the placenta. We used 100ng of RNA with RIN>3 as input for RNA library prep using the KAPA RNA HyperPrep kit with RiboErase HMR (#08098131702, Roche Diagnostics Corporation, Indianapolis, IN, USA). The kit was designed for human, mouse, and rat samples, but it also effectively reduced ribosomal RNA in our NHP RNA isolates. Libraries were prepared following the manufacturer’s standard protocol. Library quantity and quality were evaluated using Qubit Fluorometer (Thermo Fisher Scientific) and TapeStation (Agilent Technologies, Santa Clara, CA, USA). Constructed libraries were sequenced on a NovaSeq 6000 Sequencing System (#20012850, Illumina, San Diego, CA, USA), which produced 2×100 nucleotide stranded paired-end reads.

Raw sequences were aligned to the *Macaca nemestrina* reference genome from Ensembl (Mnem_1.0, INSDC Assembly GCA_000956065.1) utilizing STAR in Partek Flow, a bioinformatics software suite (Partek, Chesterfield, MO, USA). Ensembl gene symbol conversions were used to ensure consistency across all gene IDs. As the quality score was greater than 35 across all samples, no bases were trimmed, and 100% of sequencing bases overlapped with read lengths of the reference genome. The raw count matrix was downloaded from Partek. Subsequent statistical processing and analysis of count data were performed within the R statistical computing environment (R version 4.4.0). Gene counts were filtered by a row mean of 10 or greater, then normalized using edgeR to implement TMM normalization. Counts were transformed into log2 counts using voom, a function in the limma package that modifies RNA-Seq data to create a normalized count matrix.^39,40^ Differential Expression (DE) analysis compared placental samples from IAV-infected animals with those from uninfected controls. Comparisons were based on a linear model fit for each gene using limma. The criteria for all DE analyses were an absolute fold change of 2 and a p-value < 0.05. Adjusted p-values were calculated using the Benjamini-Hochberg method to reduce the false discovery rate (FDR). Gene Set Enrichment Analysis was conducted across the comparison group using fgsea with the Broad Institute Molecular Signatures database.^41^

Hierarchical clustering analysis, an unsupervised machine learning approach, was performed to group total RNA-Seq profiles based on their similarity in Euclidean distance using the hclust function in R. Ward’s linkage was used to minimize variance within each cluster. The optimal number of clusters was determined using the elbow method. The resulting hierarchy was visualized using a dendrogram. Finally, an overrepresentation analysis (ORA) was performed to determine if predefined sets of genes in Gene Ontology (GO) pathways were enriched in the cluster of differentially expressed genes (DEGs) obtained from the hierarchical clustering analysis using the clusterProfiler package in R.

### Nanostring nCounter®

To validate our results from bulk RNA sequencing, we chose a quantitative multiplex method of direct digital counting of gene transcripts using the Nanostring nCounter® Analysis Systems (Nanostring Technologies Inc., Seattle, WA, USA). RNA extracted from the chorionic villous tissues of the placenta was analyzed using the Nanostring nCounter® NHP Immunology V2 gene expression codeset (#11500276, Nanostring Technologies, Inc). RNA hybridization reactions were performed according to the manufacturer’s protocol. Probe-target RNA complexes from all reactions were processed and immobilized on nCounter® cartridges using the nCounter® Prep Station. Gene transcripts were quantified with the nCounter® Digital Analyzer (#MAN-C0035-07, Nanostring Technologies, Inc).

### Statistical Analyses

All statistical analyses were performed in RStudio (using R version 4.4.0) or nSolver (nCounter data only), both of which use the R programming language. A p-value of less than 0.05 was considered significant, and 0.05-0.1 was reported considering the modest sample size. Groups were compared using non-parametric tests (Wilcoxon rank sum) and Fisher’s exact tests. Spearman’s rank correlation coefficient was used to calculate correlations between two variables. Data visualizations were done using ggplot2, ggpmisc, and ComplexHeatmaps packages and their derivatives.^42,43^

## SUPPLEMENTARY MATERIALS

**Figs. S1 to S13.**

Fig. S1. Cytokines in Placental Chorionic Villous Tissue

Fig. S2. Cytokines in Amniotic Fluid

Fig. S3. Gestational-Age Matching of Placental Immunophenotyping

Fig. S4. Placental nCounter Gene Counts of Differentially Expressed Genes

Fig. S5. Over-Representation Analysis of Cluster 2 Differentially Expressed Genes in Figure 5A

Fig. S6. Cluster 1 Differentially Expressed Genes in Figure 5A

Fig. S7. Cluster 2 Differentially Expressed Genes in Figure 5A

Fig. S8. Correlations Between Placental Cytokines and Maternal Disease Indicators

Fig. S9. Correlations Between Placental Immunophenotyping and Maternal Disease Indicators

Fig. S10. Immunophenotyping of the Fetal Lung

Fig. S11. Immunophenotyping of the Fetal Lymph Node

Fig. S12. Immunophenotyping of Fetal Whole Blood

Fig. S13. Gestational-Age Matching of Fetal Immunophenotyping

Figure S14. Representative Gating Strategy for Granulocytes Immunophenotyping

Figure S15. Representative Gating Strategy for Immunophenotyping of Lymphoid Cells

Figure S16. Representative Gating Strategy for Monocytes Immunophenotyping

**Tables S1 to S4.**

Table S1. Animal Demographics

Table S2. Lung and Placental Pathology

Table S3. IAV H1N1 Viral Testing and Detection in Fetal Tissues

Table S4. Flow Cytometry Immunophenotyping Gates

## General

We appreciate the dedicated team of people supporting research at the Washington National Biomedical Research Center and the University of Washington, who have made this work possible. Specifically, we thank animal technicians, veterinarians, administrative personnel, Institutional Animal Care Use Committee members, and Environmental Health & Safety personnel. Without their efforts, these experiments would not have been possible.

## Author contributions

Conceptualization: KMAW, OC

Data curation: OC, SS, JM, JC

Methodology: OC, SS, HH

Investigation: OC, SS, HH, BDR, HZ, AL, AV, GM, RVA, JM, JC, RPK, ML, EL, HL, AC, MB, BC, EM, SR, BM, SW, AG, AB, CE, WMD, TBL, MNF, MM, AO, MC, LR, DHF, KMAW

Visualization: OC, SS, JM, JC

Funding acquisition: KMAW, OC, GM, DF, ML

Project administration: KMAW

Formal analysis: OC, JM (statistician), JC

Supervision: KMAW, DF

Writing – original draft: OC, KMAW, SS

Writing – review & editing: OC, KMAW, SS, RPK

## Data and materials availability

The data are openly available on several platforms. The metadata, cytokine, RT-qPCR, and flow cytometry are available in the Dryad data repository at https://doi.org/10.5061/dryad.c866t1gjf. The placenta total RNA-Seq data is available through the NIH Gene Expression Omnibus (GEO) via the GSE289117 accession number. Finally, the code used for the computational analysis and to generate the plots is available in GitHub at https://github.com/kadamswaldorflab.

