## Supplemental Information for "Influenza A Virus Infection Induces Immune Dysregulation in the Placenta and Fetus Without Vertical Transmission in Nonhuman Primates"

Table S1. Animal Demographics

| ID | Inoculum | Inoculation to Delivery Interval (Days) | Dam Age at Necropsy (Years) | Fetal Age at Inoculation (Days) | Fetal Age at Delivery (Days) | Fetal Sex | Fetal Weight(g) |
|---|---|---|---|---|---|---|---|
| CTRL1 | Media | 100 | 14.0 | 59 | 160 | F | 431.9 |
| CTRL2 | Media | 23 | 13.5 | 132 | 155 | M | 573 |
| CTRL3 | Media | 58 | 7.4 | 99 | 157 | F | 351 |
| CTRL4 | Media | 92 | 12.0 | 64 | 155 | F | 467.2 |
| CTRL5 | Media | 20 | 5.2 | 138 | 158 | F | 499.2 |
| CTRL6 | Media | 21 | 9.7 | 133 | 154 | F | 411.9 |
| CTRL7 | Media | 20 | 10.8 | 127 | 147 | M | 469 |
| CTRL8 | Media | 3 | 12.9 | 141 | 144 | M | 412.6 |
| CTRL9 | Media | 3 | 5.3 | 145 | 148 | F | 340.3 |
| CTRL10 | Saline | 7 | 10.1 | 136 | 143 | M | 405 |
| CTRL11 | Saline | 7 | 5.6 | 138 | 145 | M | 316 |
| CTRL12 | Saline | 7 | 10.7 | 132 | 139 | M | 340 |
| CTRL13 | Saline | 1 | 5.7 | 127 | 128 | M | 242 |
| CTRL14 | Saline | 1 | 6.1 | 133 | 134 | F | 264 |
| CTRL15 | Saline | 1 | 14.7 | 135 | 136 | F | 299.5 |
| CTRL16 | Saline | 1 | 12.7 | 131 | 132 | F | 298.6 |
| FLU1 | H1N1 | 5 | 12.2 | 125 | 130 | M | 291 |
| FLU2 | H1N1 | 5 | 12.7 | 126 | 131 | F | 297.2 |
| FLU3 | H1N1 | 5 | 6.6 | 127 | 132 | F | 258.3 |
| FLU4 | H1N1 | 5 | 15.5 | 138 | 143 | F | 358 |
| FLU5 | H1N1 | 5 | 10.3 | 134 | 139 | F | 372.1 |
| FLU6 | H1N1 | 5 | 7.0 | 131 | 135 | M | 334.2 |
| FLU7 | H1N1 | 5 | 13.8 | 129 | 134 | M | 318.1 |
| FLU8 | H1N1 | 5 | 8.5 | 127 | 132 | M | 277.1 |
| FLU9 | H1N1 | 5 | 7.0 | 116 | 121 | M | 264.1 |
| FLU10 | H1N1 | 5 | 4.4 | 126 | 131 | F | 316.9 |
| FLU11 | H1N1 | 5 | 11.8 | 134 | 139 | M | 356.3 |

This table shows the inoculum, inoculation to delivery interval, fetal age at inoculation and delivery, fetal sex, and fetal weight. Abbreviations: F, female; M, male

Table S2. Lung and Placental Pathology

| ID | Dam | Placenta | | | |
|---|---|---|---|---|---|
| | **Maternal Lung Pathology** | **Maternal Inflammatory Response** | **Fetal Inflammatory Response** | **Infarction** | **Necrotizing Villitis** |
| CTRL1 | None | NEG | NEG | NEG | NEG |
| CTRL2 | None | NEG | NEG | NEG | NEG |
| CTRL3 | None | NEG | NEG | NEG | NEG |
| CTRL4 | None | NEG | NEG | NEG | NEG |
| CTRL5 | None | NEG | NEG | **POS** | NEG |
| CTRL6 | None | NEG | NEG | NEG | NEG |
| CTRL7 | None | NEG | NEG | NEG | **POS** |
| CTRL8 | None | NEG | NEG | NEG | **POS** |
| CTRL9 | None | NEG | NEG | NEG | NEG |
| CTRL10 | None | NEG | NEG | NEG | NEG |
| CTRL11 | None | NEG | NEG | NEG | **POS** |
| CTRL12 | None | NEG | NEG | NEG | NEG |
| CTRL13 | None | NEG | NEG | NEG | NEG |
| CTRL14 | None | NEG | NEG | NEG | NEG |
| CTRL15 | None | NEG | NEG | NEG | **POS** |
| CTRL16 | None | NEG | NEG | NEG | NEG |
| FLU1 | Pneumonia, multifocal to coalescing, mild to moderate, mixed, with multifocal pleural fibrosis | NEG | NEG | NEG | NEG |
| FLU2 | Pneumonia, multifocal to diffuse, interstitial, mild to moderate, mixed, with multifocal pleural fibrosis | NEG | NEG | NEG | NEG |
| FLU3 | Pneumonia, interstitial to diffuse, mild (left) to severe (right), with alveolar edema and hyaline membranes | NEG | NEG | **POS** | NEG |
| FLU4 | Mild interstitial pneumonitis, mixed, with mild deep alveolar edema | NEG | NEG | NEG | **POS** |

| | | | | | |
|---|---|---|---|---|---|
| FLU5 | Pneumonia, multifocal, moderate, mixed, with edema and respiratory epithelial hyperplasia | NEG | NEG | **POS** | NEG |
| FLU6 | Pneumonia, neutrophilic to mixed, severe, with edema | NEG | NEG | NEG | **POS** |
| FLU7 | Pneumonia, interstitial, mild, chronic to mixed, with peribronchial lymphoid hyperplasia | NEG | NEG | **POS** | NEG |
| FLU8 | Pneumonia, interstitial, mild to severe, neutrophilic to mixed, with alveolar edema and fibrin | NEG | NEG | NEG | **POS** |
| FLU9 | Severe pneumonia with pleural effusion and hilar lymphadenopathy | NEG | NEG | NEG | **POS** |
| FLU10 | Pneumonia, multifocal, mild to moderate, interstitial, neutrophilic to mixed, with alveolar edema | NEG | NEG | NEG | **POS** |
| FLU11 | Severe pneumonia with pleural effusion and hilar lymphadenopathy | NEG | NEG | NEG | NEG |

This table shows a summary of maternal lung histopathology and placental pathology. Abbreviations: NEG, negative; POS, positive.

Table S3. IAV H1N1 Viral Testing and Detection in Fetal Tissues

| Fetal ID | Tissue | | | | | | | | |
|---|---|---|---|---|---|---|---|---|---|
| | **Brain** | **Brainstem** | **Spinal Cord** | **Meninges** | **Heart** | **Liver** | **Lung** | **Spleen** | **Thymus** |
| FLU1 | Neg | Neg | Neg | Neg | Neg | Neg | Neg | Neg | Neg |
| FLU2 | Neg | Neg | Neg | Neg | Neg | Neg | Neg | Neg | Neg |
| FLU3 | Neg | Neg | Neg | NT | Neg | Neg | Neg | Neg | Neg |
| FLU4 | Neg | Neg | Neg | NT | Neg | Neg | Neg | Neg | Neg |
| FLU5 | Neg | Neg | Neg | NT | Neg | Neg | Neg | Neg | Neg |
| FLU6 | Neg | NT | Neg | NT | Neg | Neg | Neg | Neg | Neg |
| FLU7 | Neg | NT | Neg | NT | NT | Neg | Neg | Neg | Neg |
| FLU8 | Neg | NT | Neg | NT | Neg | Neg | Neg | Neg | Neg |
| FLU9 | NT | NT | NT | NT | NT | NT | NT | NT | NT |
| FLU10 | Neg | Neg | Neg | NT | Neg | Neg | Neg | Neg | Neg |
| FLU11 | Neg | Neg | Neg | NT | Neg | Neg | Neg | Neg | Neg |

RT-qPCR was performed on samples from different organs of the fetuses from IAV H1N1-infected dams. FLU 9 was stillborn with severe autolysis, so the RNA quality of the tissue was insufficient for RT-qPCR testing. Abbreviations: NT, not tested; and Neg, negative.

Table S4. Flow Cytometry Immunophenotyping Gates

| **Cell Type** | **Markers** |
|---|---|
| B cells | CD14-/CD3- CD16-/HLA-DR+ CD20+ |
| CD4+ T cells | CD14-/CD16- CD3+/CD4+ CD8- |
| CD4+CD8+ T cells | CD14-/CD3- CD16+/CD4+CD8+ |
| CD8+ NK cells | CD14-/CD3- CD16+/CD8+ NKG2a+ |
| CD8+ T cells | CD14-/CD16- CD3+/CD4- CD8+ |
| Classical monocytes | CD3- CD20-/ HLA-DR+/ CD16- CD14+ |
| Intermediate monocytes | CD3- CD20-/ HLA-DR+/ CD16+ CD14hi |
| Myeloid dendritic cells | CD3- CD20-/ HLA-DR+/CD14-/ CD11c+ CD123- |
| NKT cells | CD14-/CD3+ CD16+ |
| Non-classical monocytes | CD3- CD20-/ HLA-DR+/ CD16+ CD14mid |
| Plasmacytoid dendritic cells | CD3- CD20-/ HLA-DR+/CD14-/ CD11c- CD123+ |

Overview of antibody gating used to define immune cell subsets of interest. All cell populations are live and are CD45+ singlets.

#### Figure S1. Cytokines in Placental Chorionic Villous Tissue

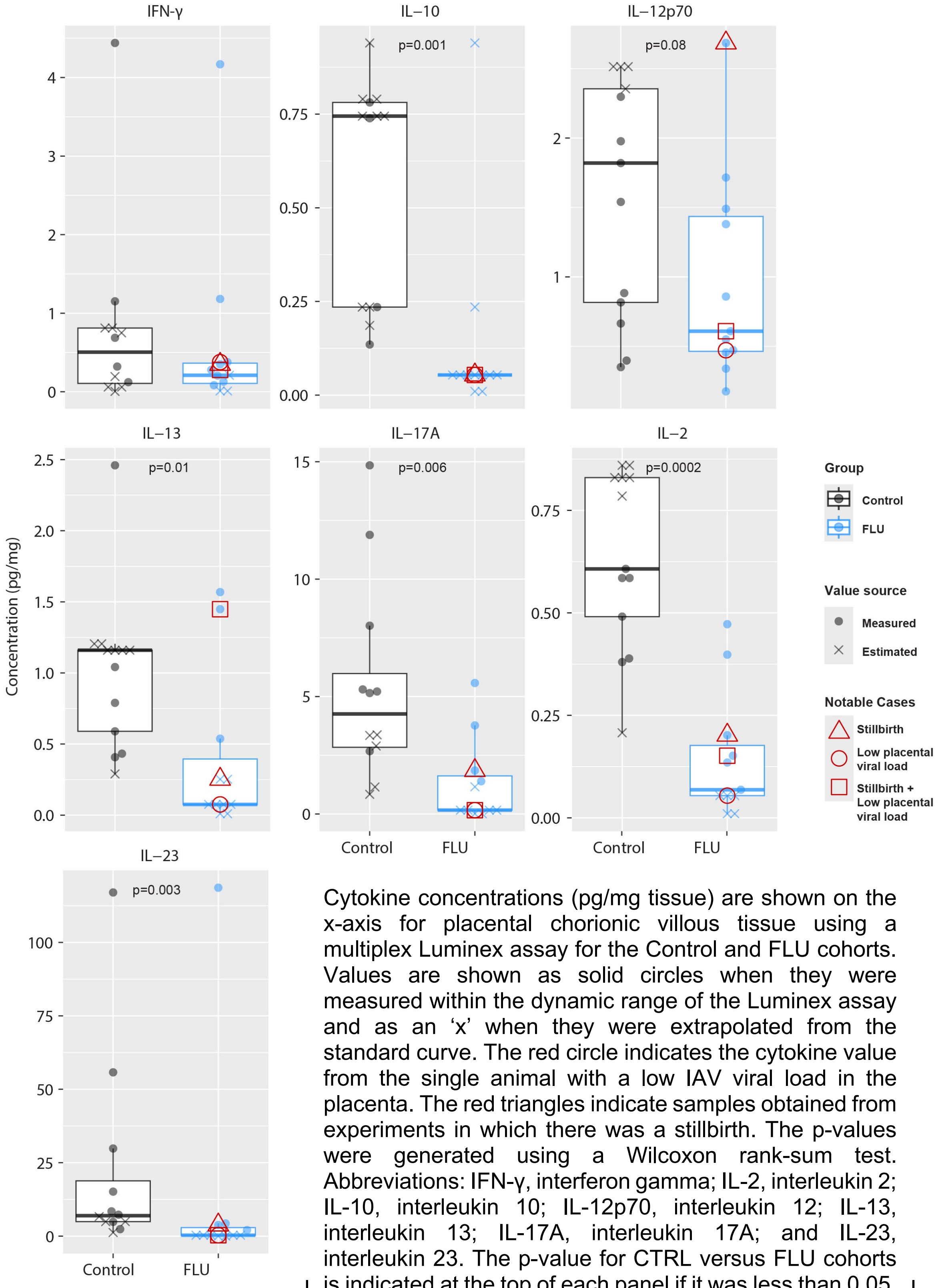

Cytokine concentrations (pg/mg tissue) are shown on the x-axis for placental chorionic villous tissue using a multiplex Luminex assay for the Control and FLU cohorts. Values are shown as solid circles when they were measured within the dynamic range of the Luminex assay and as an 'x' when they were extrapolated from the standard curve. The red circle indicates the cytokine value from the single animal with a low IAV viral load in the placenta. The red triangles indicate samples obtained from experiments in which there was a stillbirth. The p-values were generated using a Wilcoxon rank-sum test. Abbreviations: IFN-γ, interferon gamma; IL-2, interleukin 2; IL-10, interleukin 10; IL-12p70, interleukin 12; IL-13, interleukin 13; IL-17A, interleukin 17A; and IL-23, interleukin 23. The p-value for CTRL versus FLU cohorts is indicated at the top of each panel if it was less than 0.05.

#### Figure S2. Cytokines in Amniotic Fluid

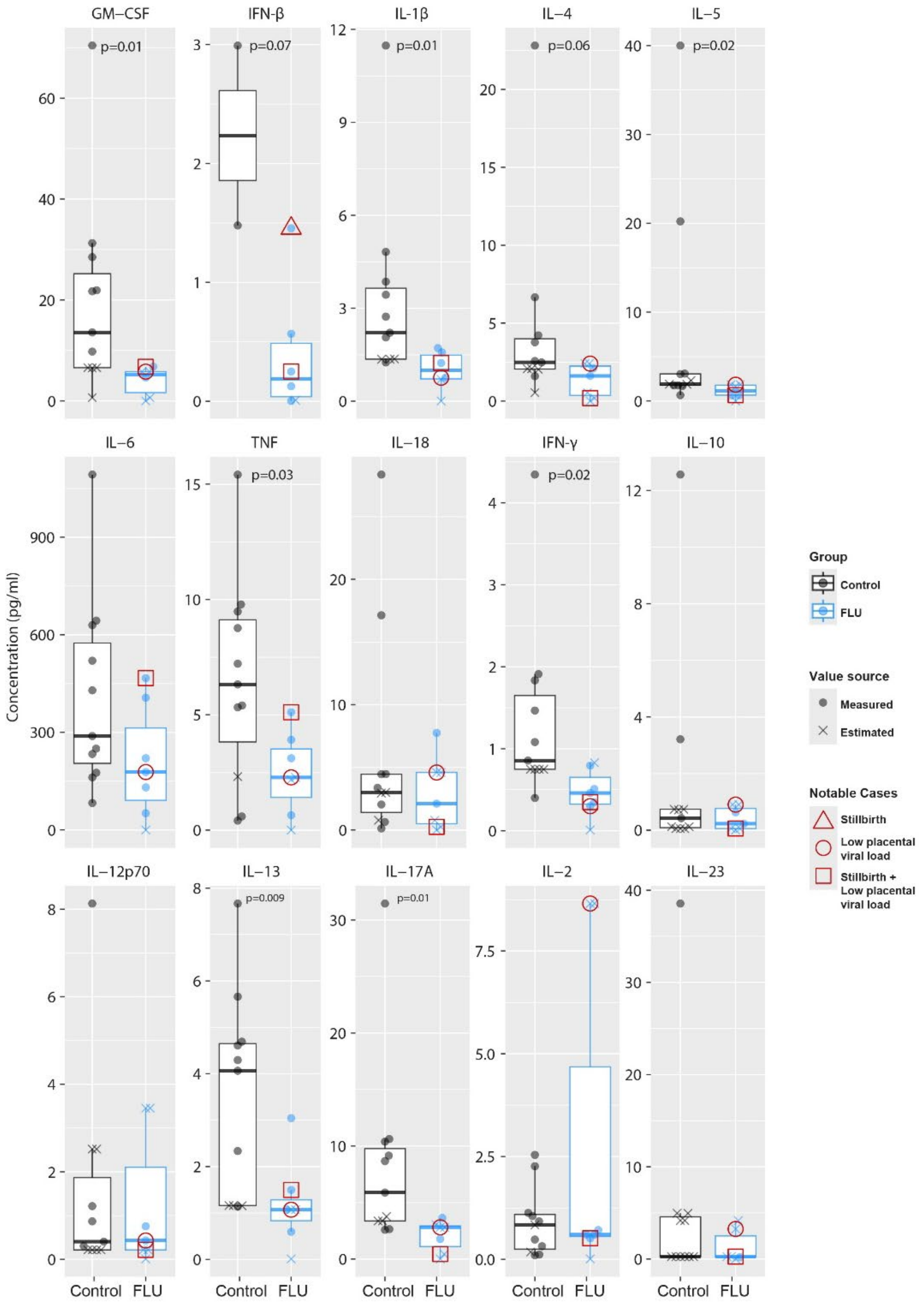

Cytokine concentrations (pg/ml) are shown on the x-axis for amniotic fluid using a multiplex Luminex assay for the Control and FLU cohorts. Values are shown as open circles when they were measured within the working range of the Luminex assay and as an 'x' when they were outside the working range and estimated by extrapolation from the standard curve. The red circle indicates the cytokine value from the single animal with a low IAV viral load in the placenta. The red triangles indicate samples obtained from experiments in which there was a stillbirth. The p-values were generated using a Wilcoxon rank-sum test. The p-value for CTRL vs FLU cohorts is indicated at the top of each panel if it was less than 0.05. Abbreviations: GM-CSF, granulocyte-macrophage colony-stimulating factor; IFN-β, interferon beta; IFN-γ, interferon gamma; IL-1β, interleukin 1 beta; IL-2, interleukin 2; IL-4, interleukin 4; IL-5, interleukin 5; IL-6, interleukin 6; IL-10, interleukin 10; IL-12p70, interleukin 12; IL-13, interleukin 13; IL-17A, interleukin 17A; IL-18, interleukin 18; IL-23, interleukin 23; and TNF, tumor necrosis factor.

Figure S3. Gestational-Age Matching of Placental Immunophenotyping

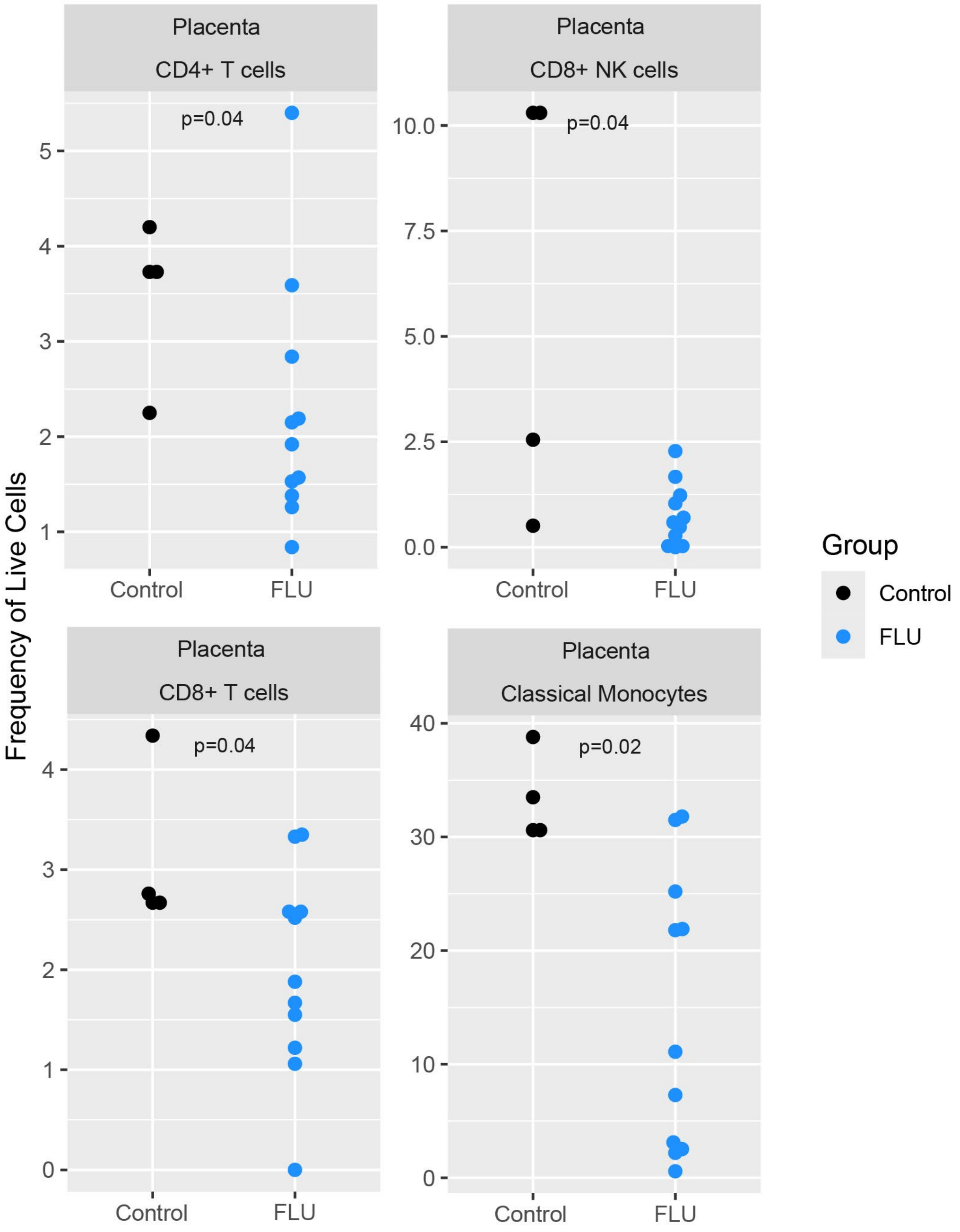

This figure illustrates the frequency of live cell populations (y-axis) of immune cell populations within the placenta chorionic villous tissue as determined by immunophenotyping (flow cytometry). Only CTRL animals with gestational age +/- 1 week of the average FLU gestational age are included. The p-values were generated using a Wilcoxon rank-sum test. The p-value for CTRL vs FLU cohorts is indicated at the top of each panel if it was found to be less than 0.1.

Figure S4. Placental nCounter Gene Counts of Differentially Expressed Genes

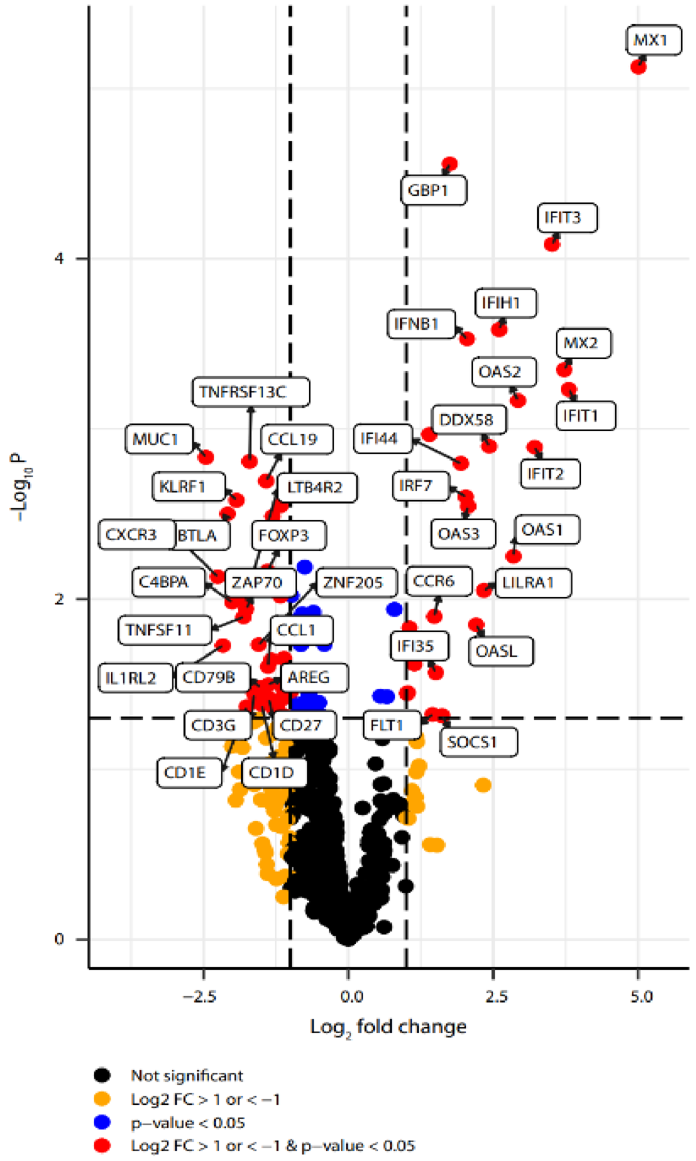

A volcano scatter plot shows the relationship between the log2 fold change and statistical significance (p-value) of differentially expressed genes between the FLU and CTRL cohorts, as determined by nCounter digital gene counting. Dot colors indicated significance and magnitude: red for genes with a significant |log2 fold change| ≥1; blue for genes with a |log2 fold change| <1 but p <0.05; gold for non-significant genes with a |log2 fold change| ≥1; and black for non-significant genes with a |log2 fold change| <1.

Figure S5. Over-Representation Analysis of Cluster 2 Differentially Expressed Genes in Figure 5A

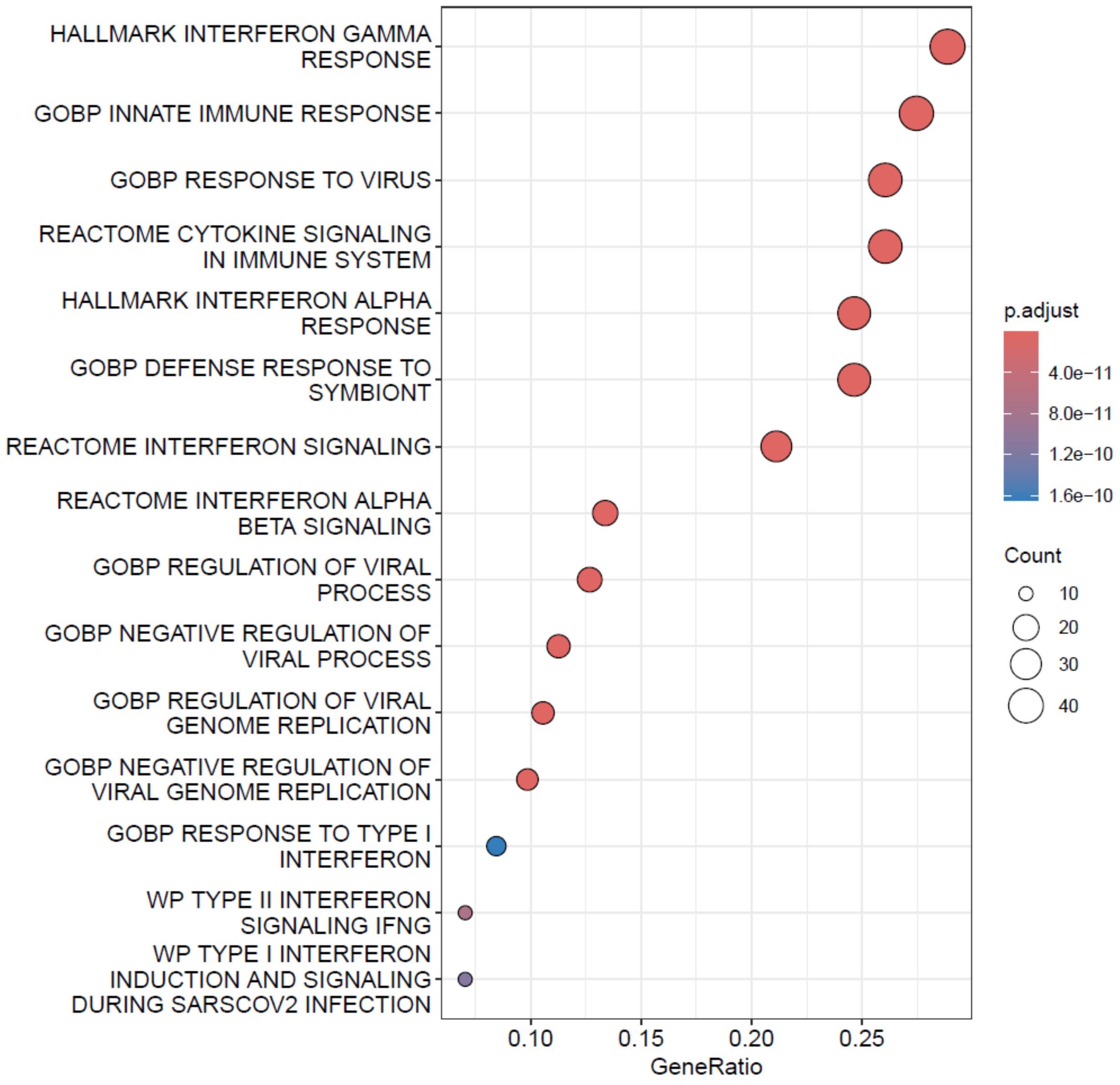

This is a bubble plot showing significantly enriched pathways from gene set over-representation analysis. Each row represents a pathway from either Gene Ontology Biological Process (GOBP), Reactome, Hallmark, or WikiPathways databases. The x-axis indicates the gene ratio (proportion of input genes associated with each pathway), while bubble size reflects the number of genes contributing to the enrichment. Bubble color corresponds to adjusted p values, with warmer colors denoting higher statistical significance. Enriched pathways include strong signatures of Type I and type II interferon signaling, viral genome regulation, and antiviral defense, highlighting the robust activation of immune responses.

Figure S6. Cluster 1 Differentially Expressed Genes in Figure 5A

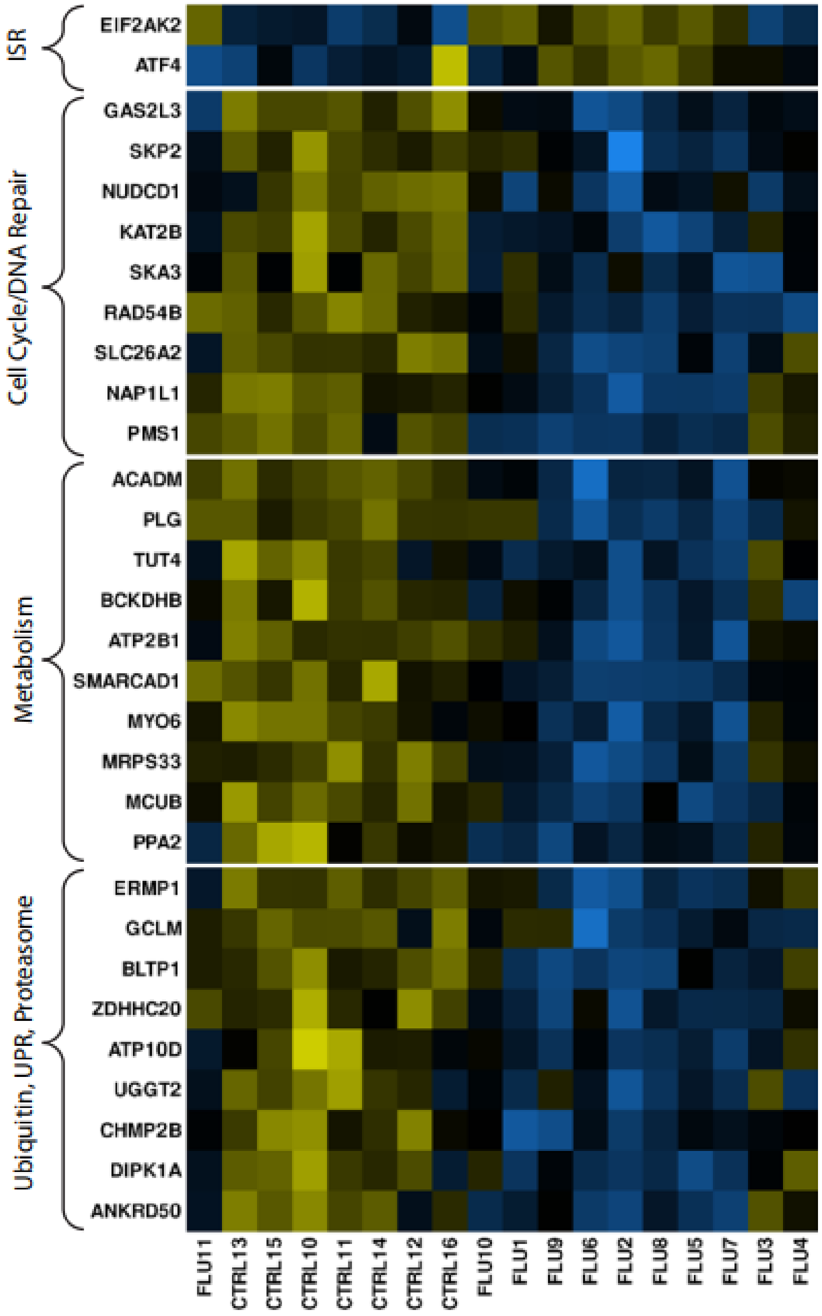

This is a heatmap showing a select set of differentially expressed genes found in Cluster 1 from the Figure 5A heatmap. The data represents normalized counts from the placental total RNA-Seq data from the FLU and CTRL cohorts. Each column represents an individual placental sample, and each row represents an individual gene. The yellow-blue color intensity indicates the gene z-score which was calculated based on the mean expression level of all animals for each gene. These genes have functions impacted by the viral-associated ISR, such as cell cycle/DNA repair, metabolism, ubiquitination, unfolded protein response (UPR), and proteasome activity. The viral-associated ISR activator (*EIF2AK2*) and mediator (*ATF4*) are shown at the top.

Figure S7. Cluster 2 Differentially Expressed Genes in Figure 5A

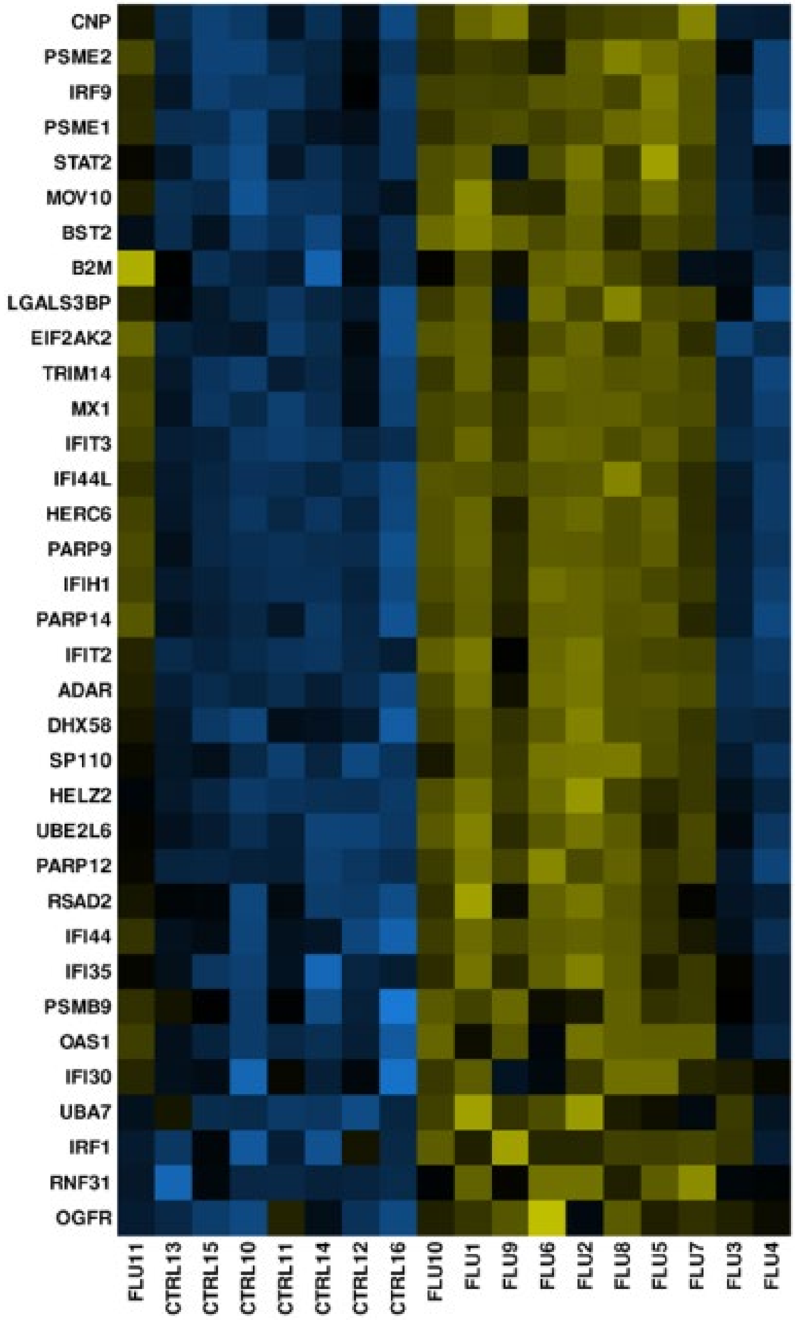

This is a heatmap showing a set of differentially expressed genes found in Cluster 2 from the Figure 5A heatmap. The data represents normalized counts from the placental total RNA-Seq data from the FLU and CTRL cohorts. Each column represents an individual placental sample, and each row represents an individual gene. The yellow-blue color intensity indicates the gene z-score which was calculated based on the mean expression level of all animals for each gene. An over-representation analysis (ORA) was performed on cluster 2 genes from panel A. Rows are genes in the Molecular Signatures Database Gene Set: Hallmark Interferon Alpha Response.

Figure S8. Correlations Between Placental Cytokines and Maternal Disease Indicators

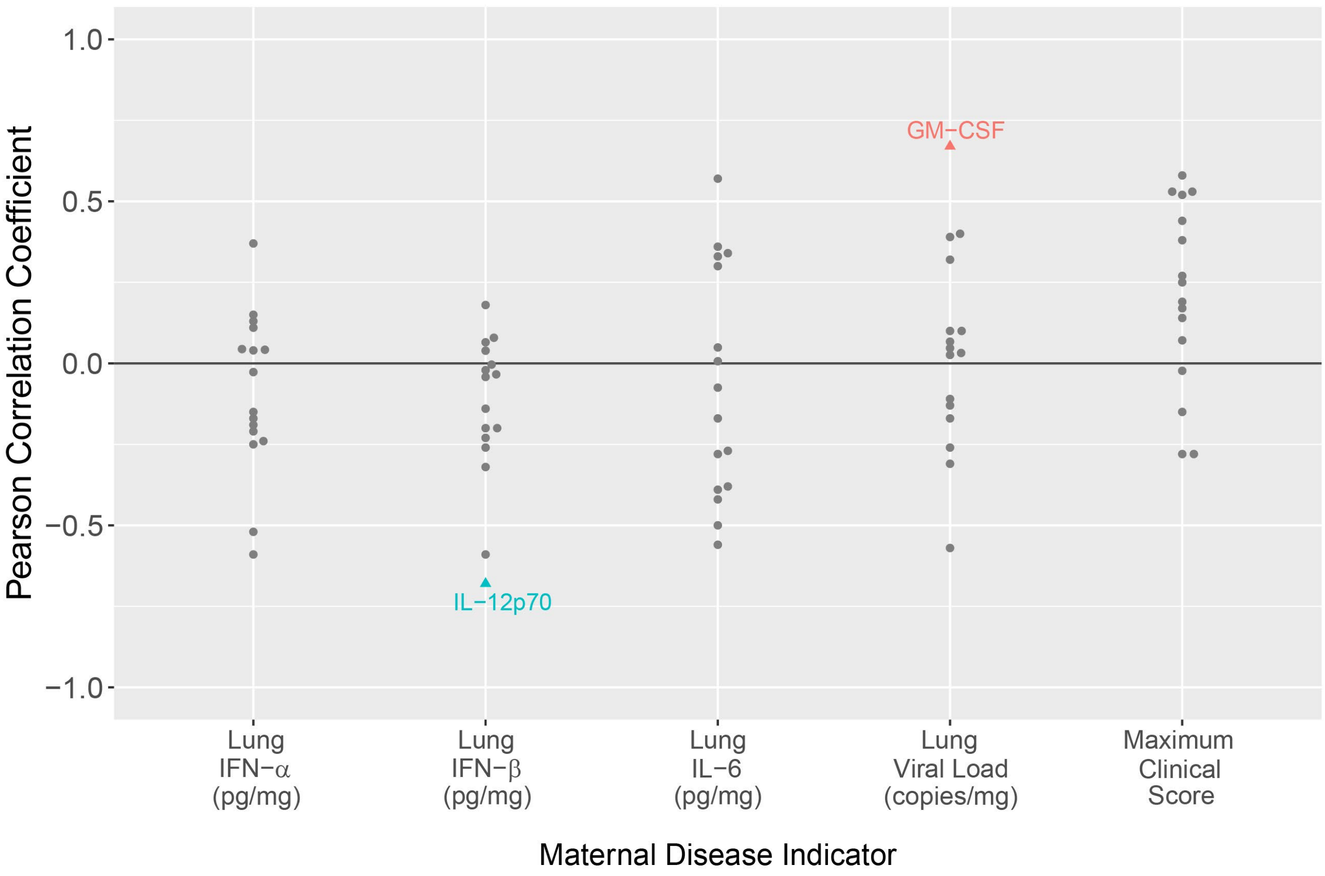

A Pearson correlation analysis was performed between placental cytokine concentrations with metrics of maternal IAV disease: lung IFN-α concentration, lung IFN-β concentration, lung IL-6 concentration, maximum clinical score, and lung viral load. Cytokine concentrations and viral load values were log10-transformed before running the correlation. The correlation coefficient, R, was plotted. Correlations with a p-value less than 0.5 are shown with a colored triangle, and correlations with a p-value greater than 0.5 are shown with a black dot. The names of cytokines with significant correlations are indicated.

Figure S9. Correlations Between Placental Immunophenotyping and Maternal Disease Indicators

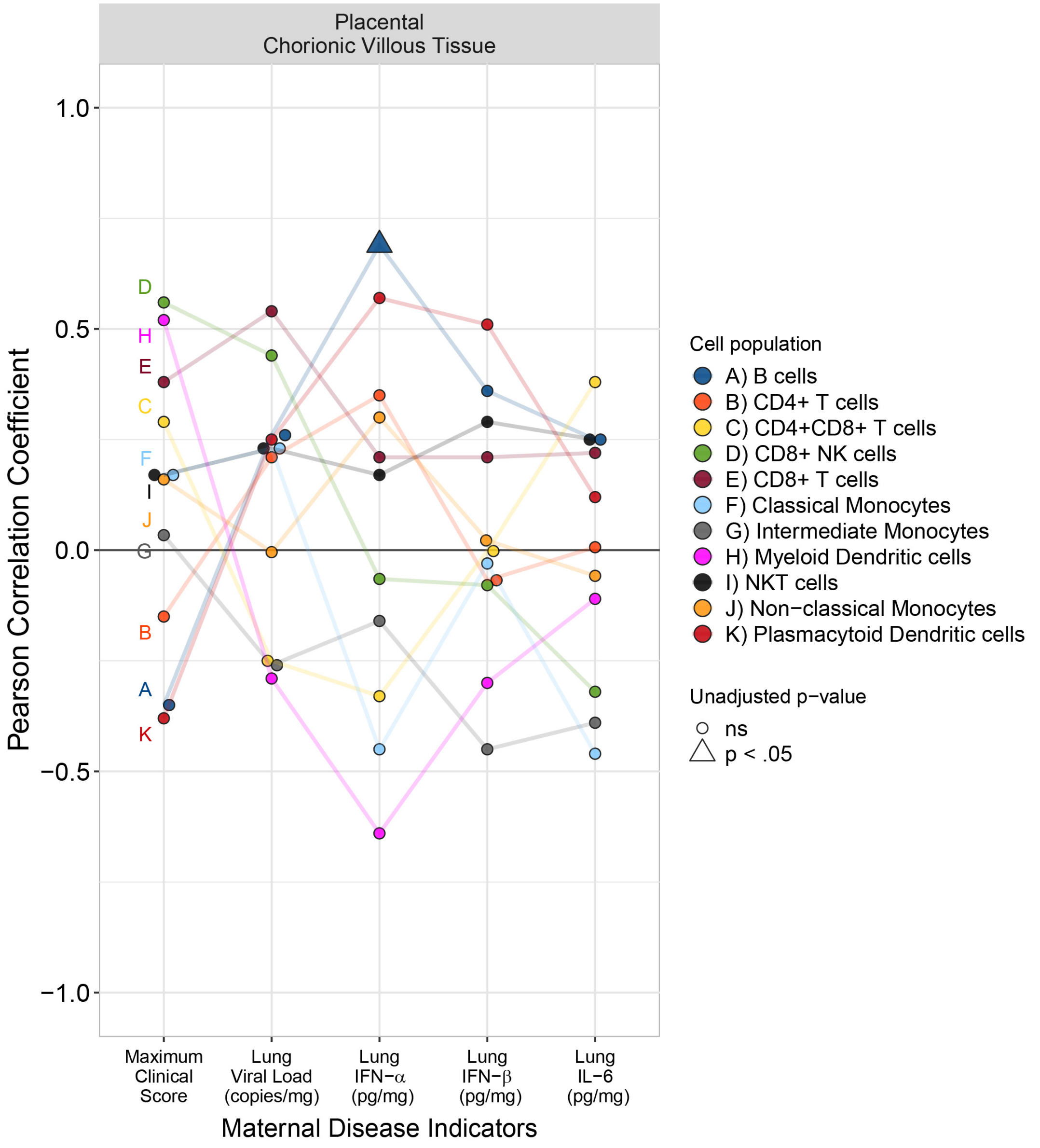

Pearson correlation analysis between maternal disease indicators (maximum clinical score, viral load, maternal lung IFN-α, IFN-β, and IL-6) and placental immune cell populations. The Pearson correlation coefficient is shown on the y-axis, and the maternal disease indicator is shown on the x-axis. Each immune cell population is shown by color and letter to aid in distinguishing populations from one another. A line connects each Pearson correlation coefficient from the same immune cell population, allowing tracking of similarities in positive and negative coefficients. Triangles show significant unadjusted p-values.

#### Figure S10. Immunophenotyping of the Fetal Lung

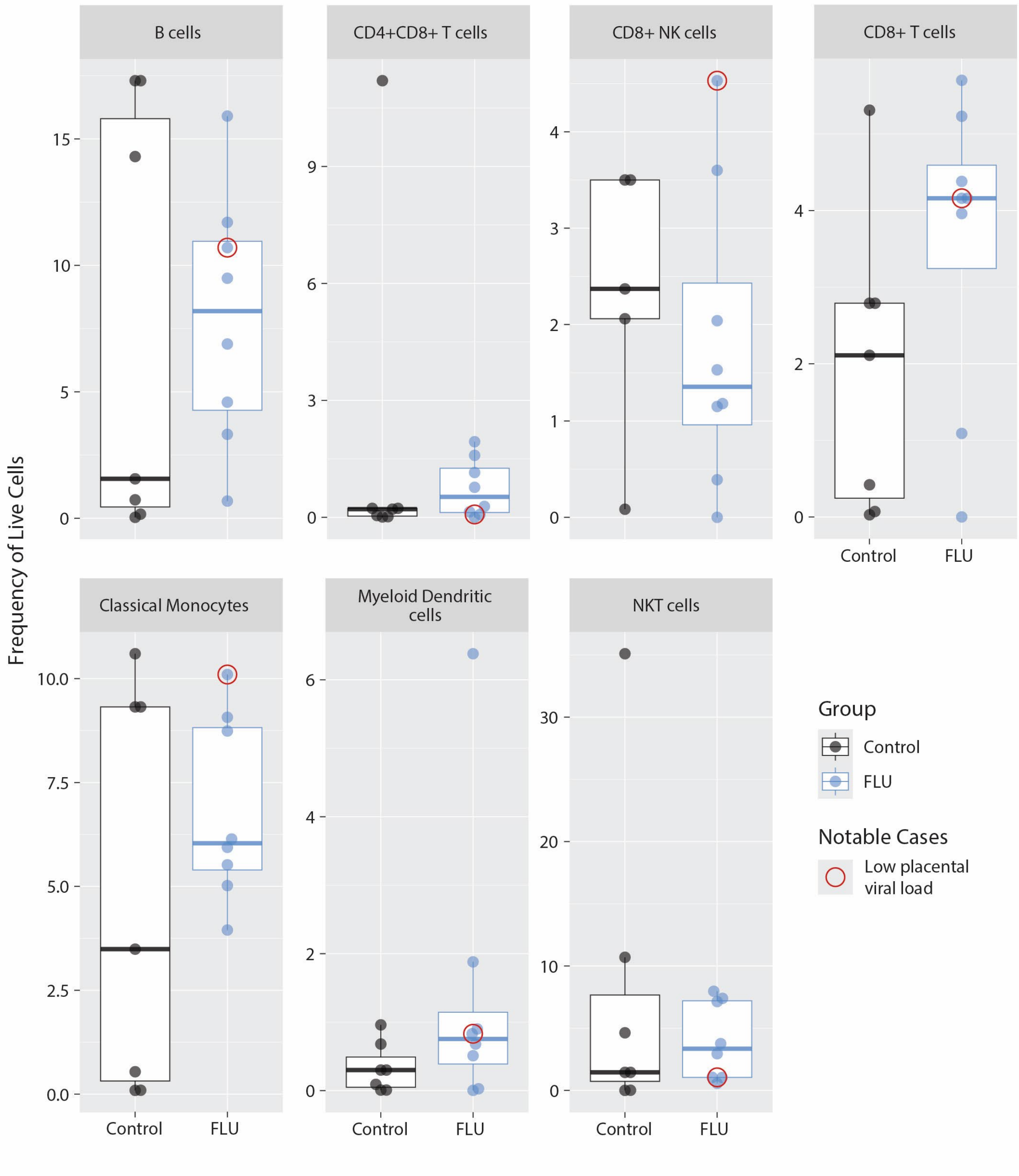

This figure illustrates the frequency of live cell populations (y-axis) of immune cell populations within the fetal lung, as determined by immunophenotyping (flow cytometry). The red circle indicates the frequency of live cells in the fetal lung from the pregnancy with a low IAV viral load in the placenta. Immunophenotyping was not done on fetal tissues from stillbirth. Abbreviations: NKT, natural killer T cells.

### Figure S11. Immunophenotyping of the Fetal Lymph Node

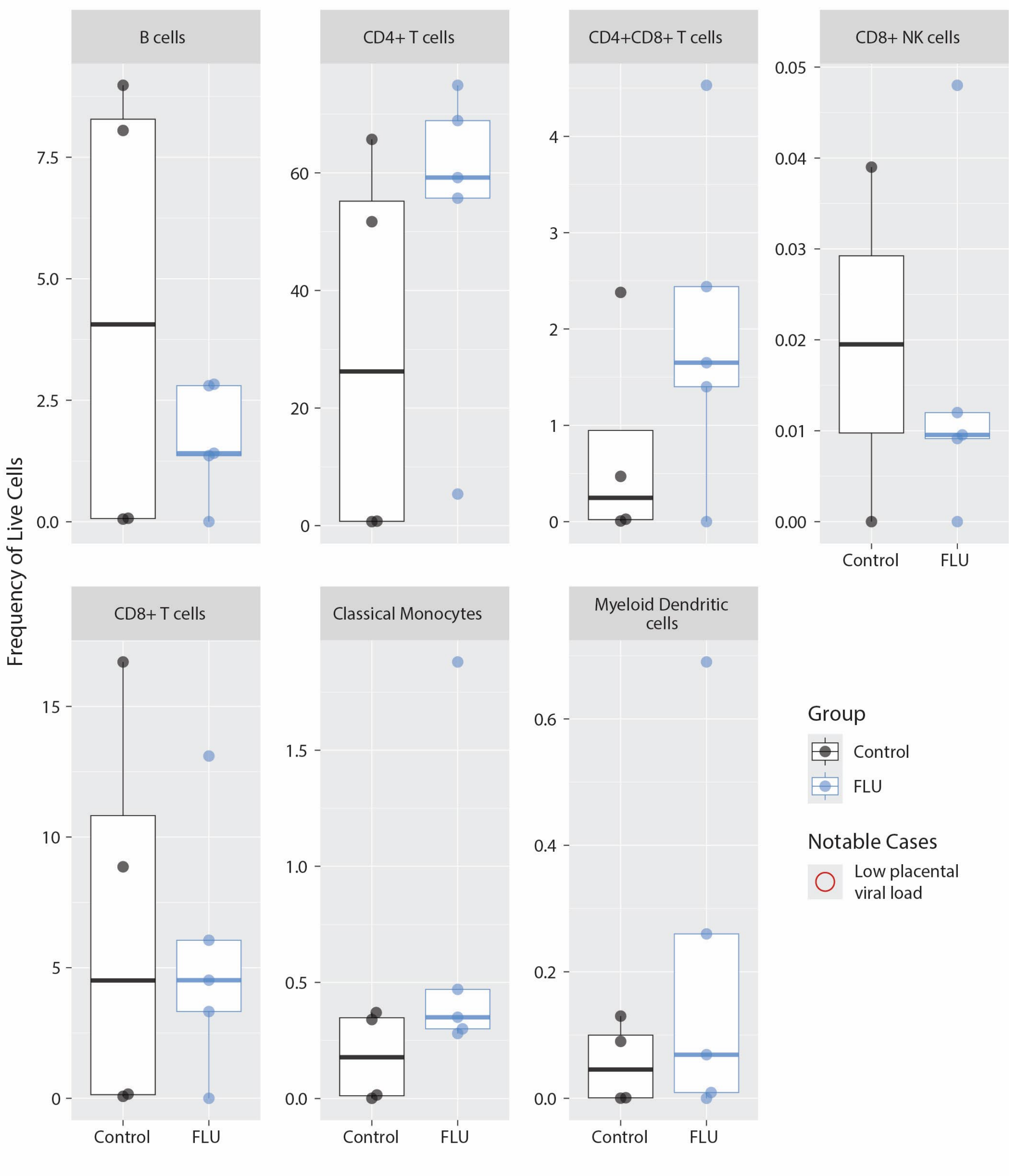

This figure illustrates the frequency of live cell populations (y-axis) of immune cell populations within the fetal mesenteric lymph nodes, as determined by immunophenotyping (flow cytometry). The red circle indicates the frequency of live cells in the fetal lung from the pregnancy with a low IAV viral load in the placenta. Immunophenotyping was not done on fetal tissues from stillbirth.

Figure S12. Immunophenotyping of Fetal Whole Blood

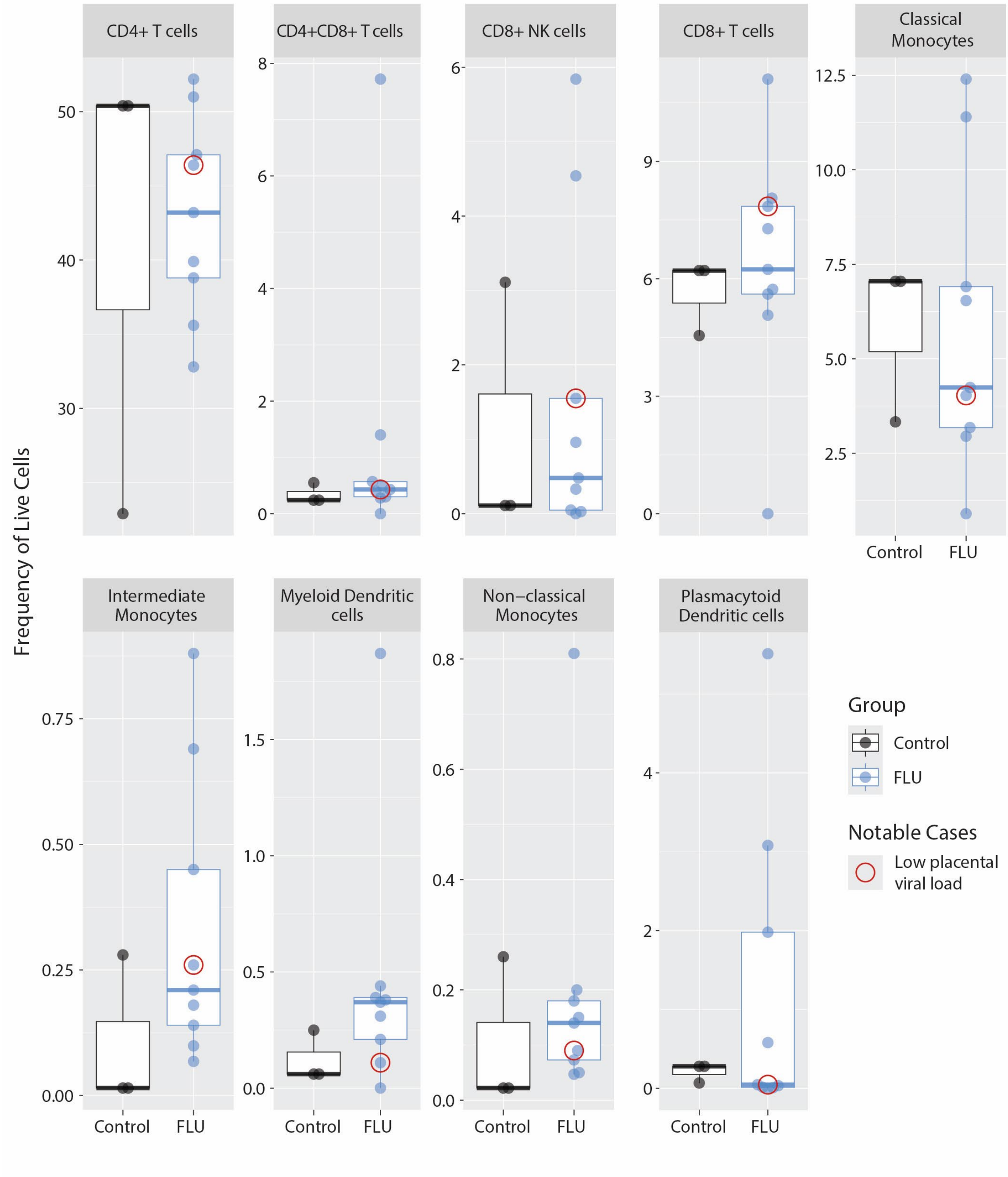

This figure illustrates the frequency of live cell populations (y-axis) of immune cell populations within the fetal whole blood, as determined by immunophenotyping (flow cytometry). The red circle indicates the frequency of live cells in the fetal lung from the pregnancy with a low IAV viral load in the placenta. Immunophenotyping was not done on fetal tissues from stillbirth.

Figure S13. Gestational-Age Matching of Fetal Immunophenotyping

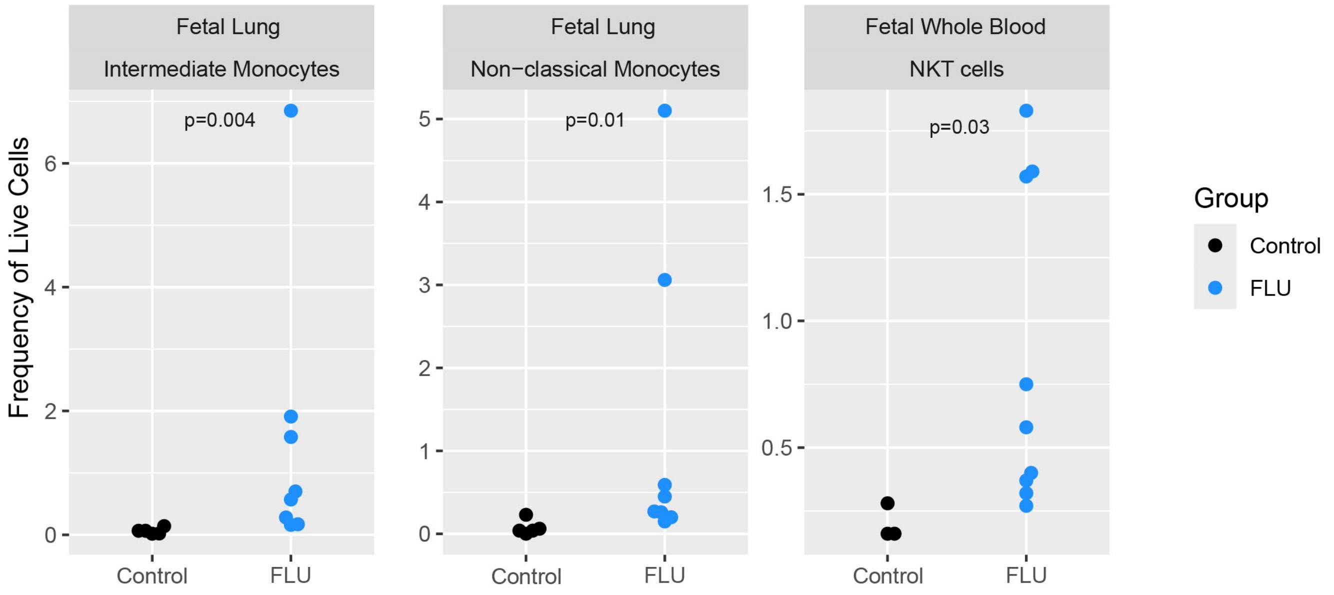

This figure illustrates the frequency of live cell populations (y-axis) of immune cell populations within the fetal lung (left and middle) and fetal whole blood (right) as determined by immunophenotyping (flow cytometry). Only CTRL animals with gestational age +/- 1 week of the average FLU gestational age are included. Immunophenotyping was not done on fetal tissues from stillbirth. The p-values were generated using a Wilcoxon rank-sum test. The p-value for CTRL vs FLU cohorts is indicated at the top of each panel if it was found to be less than 0.1.

Figure S14. Representative Gating Strategy for Granulocytes Immunophenotyping

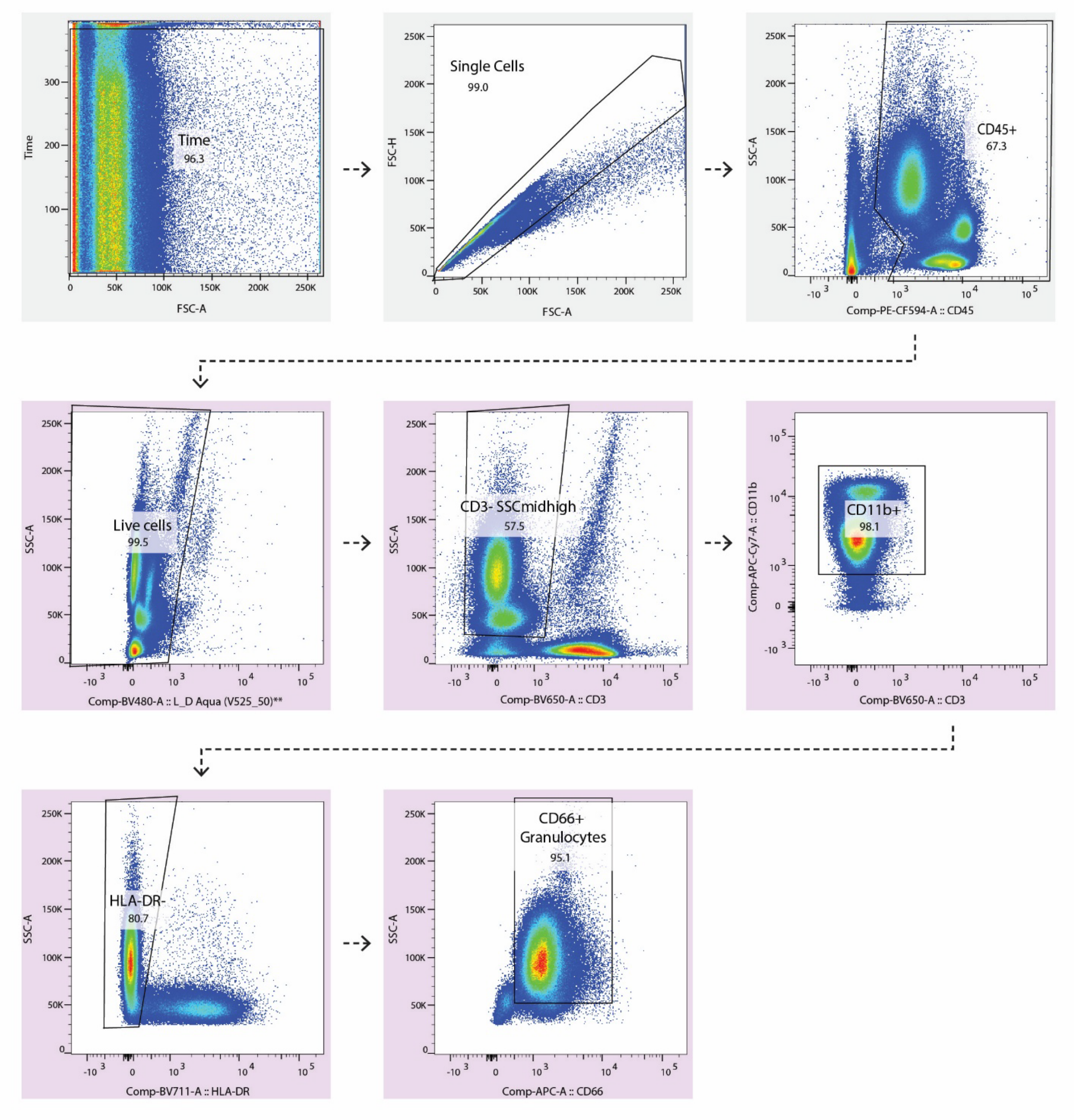

Single cells were analyzed by multiparametric flow cytometry. Doublets were excluded by forward scatter height (FSC-H) versus forward scatter area (FSC-A) gating. Leukocytes were gated by side scatter (SSC-A) and CD45 to distinguish granulocytes from non-hematopoietic events. Within CD45+ cells, viable cells were identified by exclusion of the viability dye (Aqua). Next, cells were identified as either CD3 negative, mid- and high-SSC-A cells. CD66+ granulocytes were further discriminated by gating to isolate CD11b+, HLADR-, and CD66+ cells. Shown is a representative gating scheme from one subject. The parent population name is overlayed on the plot in large text.

Figure S15. Representative Gating Strategy for Immunophenotyping of Lymphoid Cells

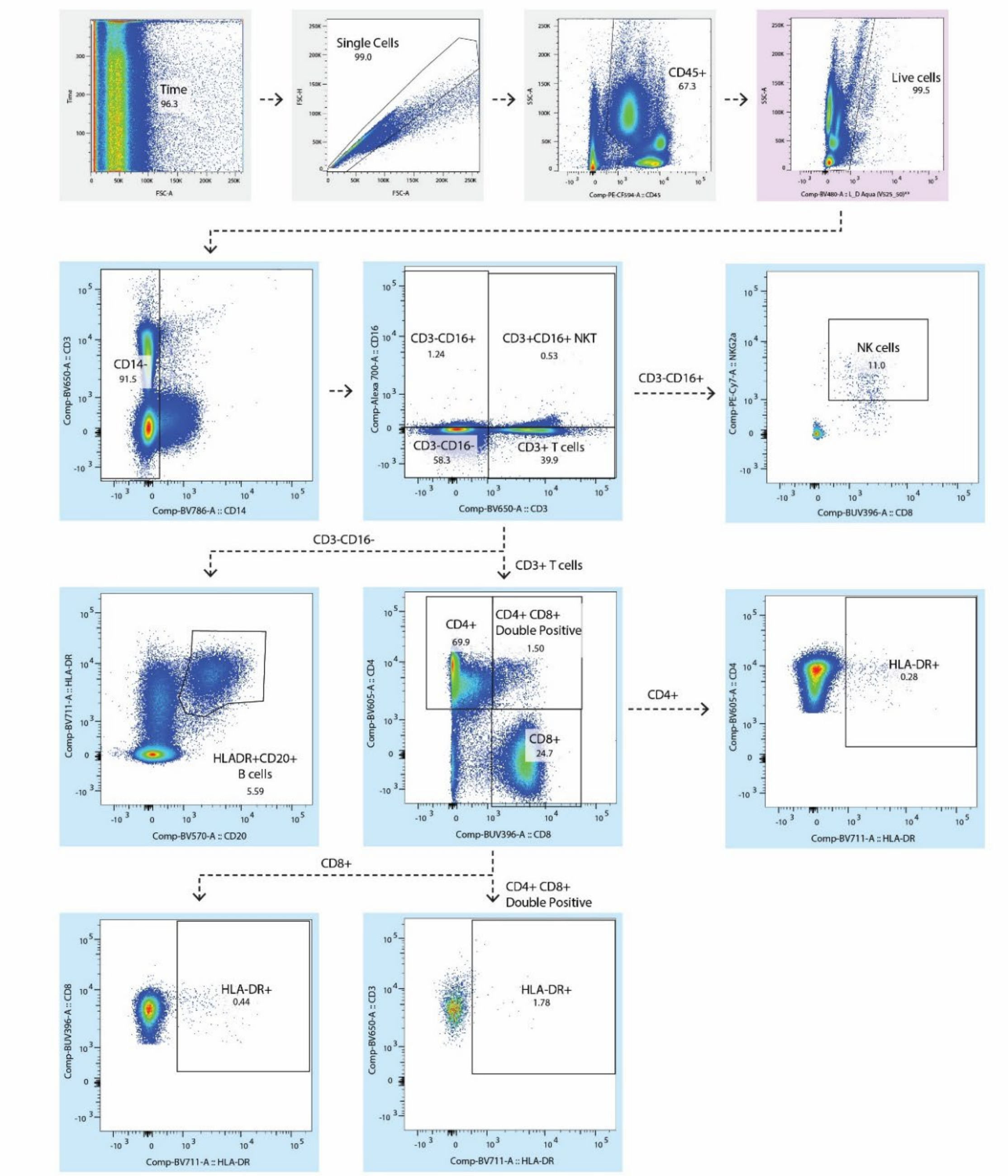

Single cells were analyzed by multiparametric flow cytometry. Doublets were excluded by forward scatter height (FSC-H) versus forward scatter area (FSC-A). Leukocytes were gated by side scatter (SSC-A) and CD45 to distinguish granulocytes from non-hematopoietic events. Within CD45+ cells, viable cells were identified by exclusion of the viability dye (Aqua). Within the lymphocyte gate, major lineages were defined using CD3, CD16, HLA-DR, and CD20. Natural killer (NK) cells were identified as CD3-/CD16+. B cells were defined as CD3-/CD16-/HLA-DR+CD20+. CD3+ T cells were subdivided into CD4+, CD8+, and double-positive (CD4+CD8+) populations. Activated T cell subsets were further characterized based on HLA-DR expression, including CD4+HLA-DR+ T cells, CD8+HLA-DR+ T cells, and CD4+CD8+HLA-DR+ T cells. Shown is a representative gating scheme from one subject. The parent population name is overlaid on the plot or next to its corresponding arrow in large text.

Figure S16. Representative Gating Strategy for Monocyte Immunophenotyping

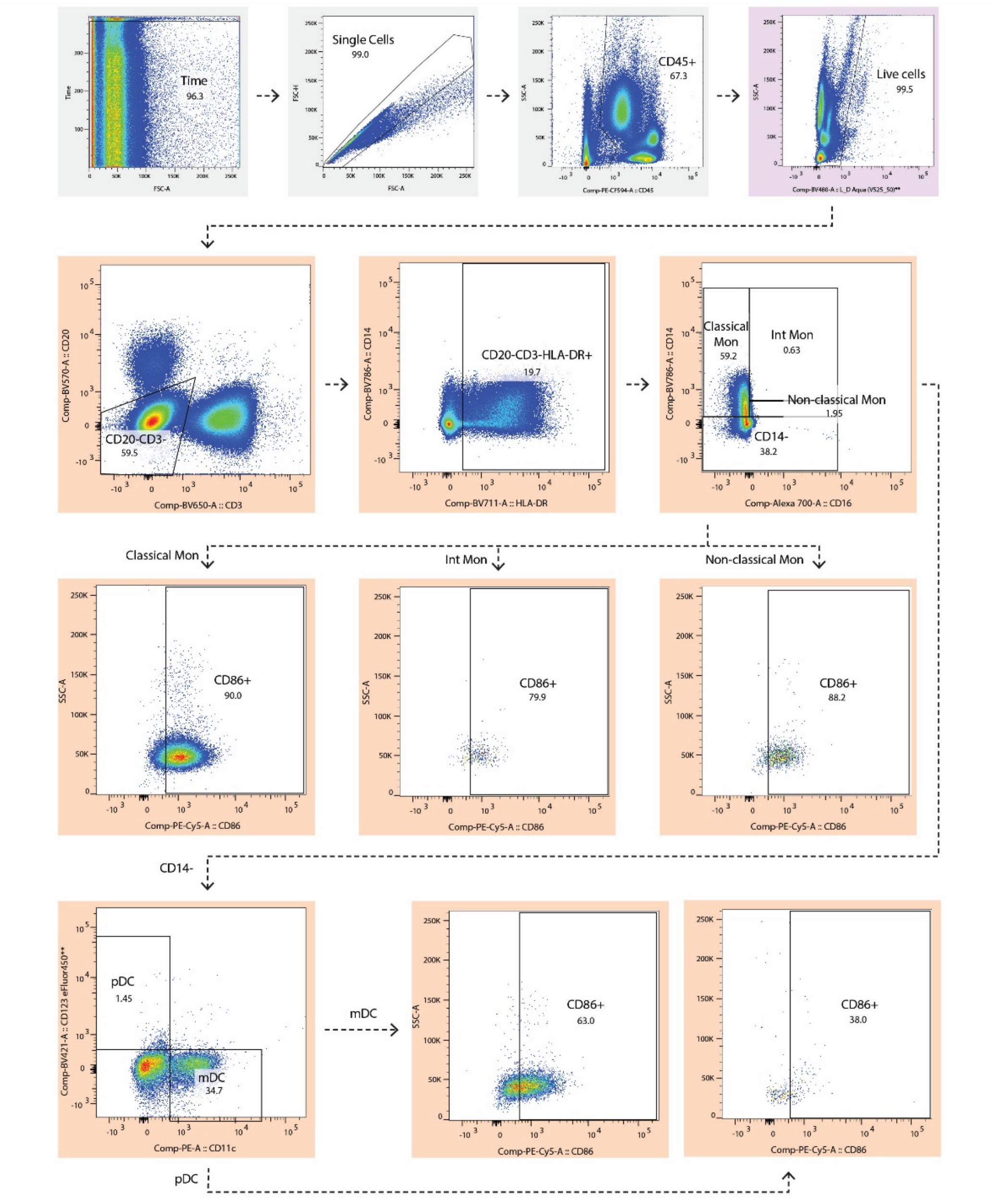

Single cells were analyzed by multiparametric flow cytometry. Doublets were excluded by forward scatter height (FSC-H) versus forward scatter area (FSC-A). Leukocytes were gated by side scatter (SSC-A) and CD45 to distinguish granulocytes from non-hematopoietic events. Within CD45+ cells, viable cells were identified by exclusion of the viability dye (Aqua). Monocytes and dendritic cells were gated from CD45+ leukocytes to select CD20-CD3- cells. (D) Within the monocyte population, subsets were defined using CD14, CD16, and CD86 expression: classical monocytes (CD14++CD16-CD86+), intermediate monocytes (CD14+CD16+CD86+), and non-classical monocytes ($CD14^{lo}$CD16++CD86+). (E) Dendritic cells were identified by exclusion of lineage markers and further subdivided based on CD11c, CD123, and CD86 expression: myeloid dendritic cells (CD11c+CD123-CD86+) and plasmacytoid dendritic cells (CD11c-CD123+CD86+). Shown is a representative gating scheme from one subject. The parent population name is overlaid on the plot or next to its corresponding arrow in large text.
